# A cryptic transcription factor regulates *Caulobacter* adhesin development

**DOI:** 10.1101/2022.05.23.493133

**Authors:** Maeve McLaughlin, David M. Hershey, Leila M. Reyes Ruiz, Aretha Fiebig, Sean Crosson

**Affiliations:** Department of Microbiology and Molecular Genetics, Michigan State University, East Lansing, MI, USA; Department of Bacteriology, University of Wisconsin, Madison, WI, USA; Department of Microbiology and Immunology, University of North Carolina, Chapel Hill, NC, USA

## Abstract

*Alphaproteobacteria* commonly produce an adhesin that is anchored to the exterior of the envelope at one cell pole. In *Caulobacter crescentus* this adhesin, known as the holdfast, facilitates attachment to solid surfaces and cell partitioning to air-liquid interfaces. An ensemble of two-component signal transduction (TCS) proteins controls *C. crescentus* holdfast biogenesis by indirectly regulating expression of HfiA, a potent inhibitor of holdfast synthesis. We performed a genetic selection to discover direct *hfiA* regulators that function downstream of the adhesion TCS system and identified *rtrC*, a hypothetical gene. *rtrC* transcription is directly activated by the adhesion TCS regulator, SpdR. Though its primary structure bears no resemblance to any defined protein family, RtrC binds and regulates dozens of sites on the *C. crescentus* chromosome via a pseudo-palindromic sequence. Among these binding sites is the *hfiA* promoter, where RtrC functions to directly repress transcription and thereby activate holdfast development. Either RtrC or SpdR can directly activate transcription of a second *hfiA* repressor, *rtrB*. Thus, environmental regulation of *hfiA* transcription by the adhesion TCS system is subject to control by an OR-gated type I coherent feedforward loop; these regulatory motifs are known to buffer gene expression against fluctuations in regulating signals. We have further assessed the functional role of *rtrC* in holdfast-dependent processes, including surface adherence to a cellulosic substrate and formation of pellicle biofilms at air-liquid interfaces. Strains harboring insertional mutations in *rtrC* have a diminished adhesion profile in a competitive cheesecloth binding assay and a reduced capacity to colonize pellicle biofilms in select media conditions. Our results add to an emerging understanding of the regulatory topology and molecular components of a complex bacterial cell adhesion control system.

**Author Summary:** A complex structure known as the envelope separates the controlled interior of bacterial cells from the external environment. The envelope regulates molecular traffic in and out of the cell and mediates physical contact with the cell’s surroundings. Bacteria often anchor specialized polymers to the exterior of their envelopes, which enable attachment to surfaces and facilitate the development of multicellular communities known as biofilms. We have discovered that an uncharacterized hypothetical gene, present in common soil and aquatic bacteria, functions to control development of a surface adhesin known as the holdfast. This gene, which we have named *rtrC*, encodes a DNA-binding protein that regulates the expression of dozens of genes in *Caulobacter*. The expression of *rtrC* results in potent activation of holdfast biosynthesis, and loss of *rtrC* results in defects holdfast-dependent processes in *Caulobacter* including the ability to colonize biofilms at the surface of water. The results presented in this study illuminate the molecular function of previously hypothetical gene, and inform understanding of the molecular processes and pathways that control bacterial adhesion and biofilm development.

## Introduction

The ability of microbial cells to adhere to surfaces and form biofilms is often a key determinant of fitness in both clinical and non-clinical contexts [1-3]. Colonization of substrates can support energy production [4], protect cells from toxic compounds [5, 6], and shield cells from grazing protist predators [7]. However, competition for resources in a multicellular biofilm can also slow growth; thus, there are evolutionary tradeoffs between surface attached and planktonic lifestyles [8]. Given that the fitness benefit of surface attachment varies as a function of environmental conditions, it follows that the cellular decision to adhere to a substrate is highly regulated.

Gram-negative bacteria of the genus *Caulobacter* are common in aquatic and soil ecosystems [9] and are dominant members of mixed biofilm communities in freshwater [10]. *Caulobacter* spp. often produce a secreted polar adhesin known as the holdfast, which enables high-affinity attachment to surfaces [11] and robust biofilm formation [12]. In the model *Caulobacter* species, *C. crescentus*, holdfast development is regulated at many levels. The transcription of holdfast synthesis genes exhibits periodic changes across the cell cycle, consistent with the developmental regulation of holdfast synthesis [13, 14]. In addition, the small protein, HfiA, is a potent post-translational inhibitor of holdfast synthesis that itself is controlled by cell cycle and environmental signals [15-17]. Holdfast biogenesis is also influenced by mechanical cues [18-20], while the second messenger cyclic-di-GMP affects both synthesis [19] and physical properties of [21] the holdfast. Additionally, an elaborate regulatory pathway comprised of multiple two-component signaling (TCS) proteins and one-component regulators controls holdfast development and surface attachment [16]. We have previously shown that a *C. crescentus* strain expressing a non-phosphorylatable allele of the *lovK* sensor histidine kinase (*lovK*_H180A_) overproduces holdfast and, consequently, has an enhanced adhesion phenotype in a biofilm assay. The *lovK*_H180A_ adhesion phenotype requires the presence of *spdS-spdR* two-component system genes and the hybrid histidine kinase *skaH* gene [16]. Two XRE-family transcription factors, RtrA and RtrB, function downstream of the TCS regulators to promote holdfast synthesis by directly repressing transcription of the holdfast inhibitor, *hfiA* (Figure 1A). Though *rtrA* and *rtrB* clearly contribute to holdfast regulation downstream of the adhesion TCS proteins, we hypothesized that there were additional regulators of *C. crescentus* holdfast biosynthesis in this pathway. Our hypothesis is based on the observation that deletion of both *rtrA* and *rtrB* does not completely abrogate holdfast synthesis when the TCS pathway is constitutively activated [16].

**Figure 1.**
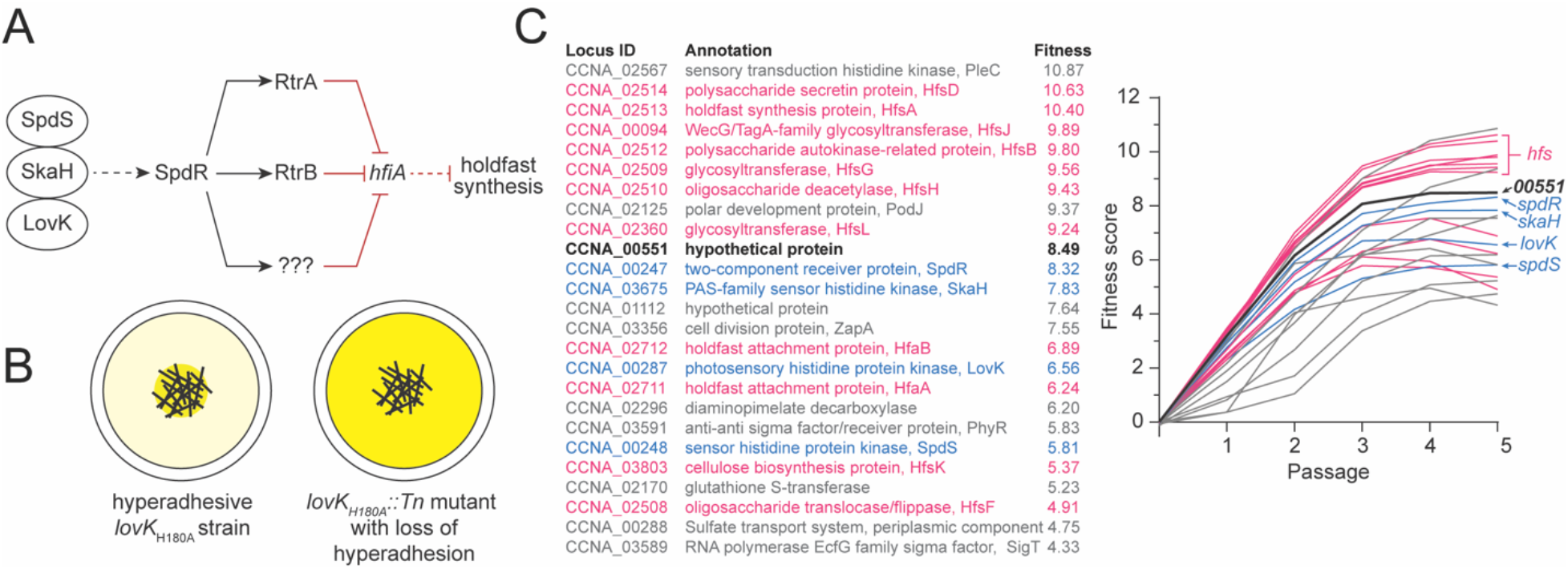
Adhesion TCS pathway and adhesion profiling of *lovK*_H180A_::*Tn* mutants. **A)** Schematic of the LovK-SpdSR-SkaH adhesion TCS system that regulates holdfast synthesis as described by Reyes-Ruiz et al. [16]. Question marks indicate postulated additional regulator(s) in the adhesion control pathway. Dashed lines indicate post-transcriptional regulation and solid lines indicate transcriptional regulation. Black arrows indicate activation and red bar-ended lines indicate repression. **B)** Genetic selection to identify insertions that disrupt the hyper-holdfast phenotype of *lovK*_H180A_. Tn-*himar* strains were cultivated and serially passaged for five days in the presence of cheesecloth (black cross-hatched lines in the center of the well). Mutants that do not permanently adhere to cheesecloth are increasingly enriched in the supernatant with each passage. Darker yellow color indicates non-adhesive *lovK*_H180A_::Tn strains that are enriched after five days of serial passaging. **C)** (left) List of the 25 genes for which transposon insertion has the largest disruptive effect on adhesion of the *lovK*_H180A_ strain. (right) Enrichment in the supernatant is reflected in an increasing calculated fitness score with each daily passage. Mutations that disrupt *lovK*_H180A_ adhesion to cheesecloth include the expected holdfast synthesis and modification genes (pink), and genes encoding the LovK-SpdSR-SkaH regulatory system (blue). The hypothetical gene *CCNA_00551* is listed in black; all remaining genes are colored grey.

To search for these postulated downstream regulators, we used a transposon sequencing approach to select for insertions that attenuate the hyper-holdfast phenotype of a *lovK*_H180A_ mutant. Our selection uncovered a gene encoding a hypothetical protein that we have named RtrC, which functions as both a transcriptional activator and repressor in *C. crescentus*. RtrC binds a pseudo-palindromic DNA motif *in vivo* and *in vitro* and activates holdfast synthesis downstream of the *lovK-spdSR-skaH* TCS ensemble by directly repressing transcription of the holdfast inhibitor, *hfiA*. RtrC, along with the response regulator SpdR, and the transcription factor RtrB form an OR-gated type I coherent feedforward loop (C1-FFL) that regulates *hfiA* transcription. C1-FFL motifs are known to buffer gene expression against transient loss of regulating signals, which often occurs in fluctuating natural environments. Beyond *hfiA*, RtrC can also directly control the transcription of dozens of other genes in *C. crescentus* via its pseudo-palindromic binding site, including genes that impact flagellar motility, cyclic-di-GMP signaling, and aerobic respiration.

## Results

### A hypothetical protein functions downstream of the TCS regulators, LovK and SpdR, to activate holdfast synthesis

An ensemble of two-component signal transduction (TCS) proteins in *C. crescentus*, including LovK and SpdR, can control holdfast synthesis by indirectly regulating transcription of *hfiA*. Two XRE-family transcription factors, RtrA and RtrB, function downstream of this TCS system to directly repress *hfiA* and thereby activate holdfast synthesis [16] (Figure 1A). However, deleting *rtrA, rtrB*, or both (as shown in [16]) has only modest effects on holdfast synthesis when the TCS system is constitutively activated (Figure 2B). We therefore reasoned that there are additional downstream regulators in this pathway that can activate *C. crescentus* holdfast synthesis. To identify genes downstream of *lovK* that regulate holdfast synthesis in both an *spdR*-dependent and *spdR*-independent manner, we constructed a randomly barcoded transposon mutant library in a *lovK* mutant background (*lovK*_H180A_) in which holdfast synthesis is constitutively activated. This barcoded library was cultivated and serially passaged in the presence of cheesecloth, a process that titrates adhesive cells from liquid medium as recently described [22]. Non-adhesive mutants become enriched in the media supernatant surrounding the cheesecloth, which is reflected as a positive fitness score when the total barcoded population is quantified (Figure 1B). Using this approach, we aimed to identify transposon insertions that ablated the hyper-holdfast phenotype of a mutant in which the adhesion pathway is constitutively active.

**Figure 2.**
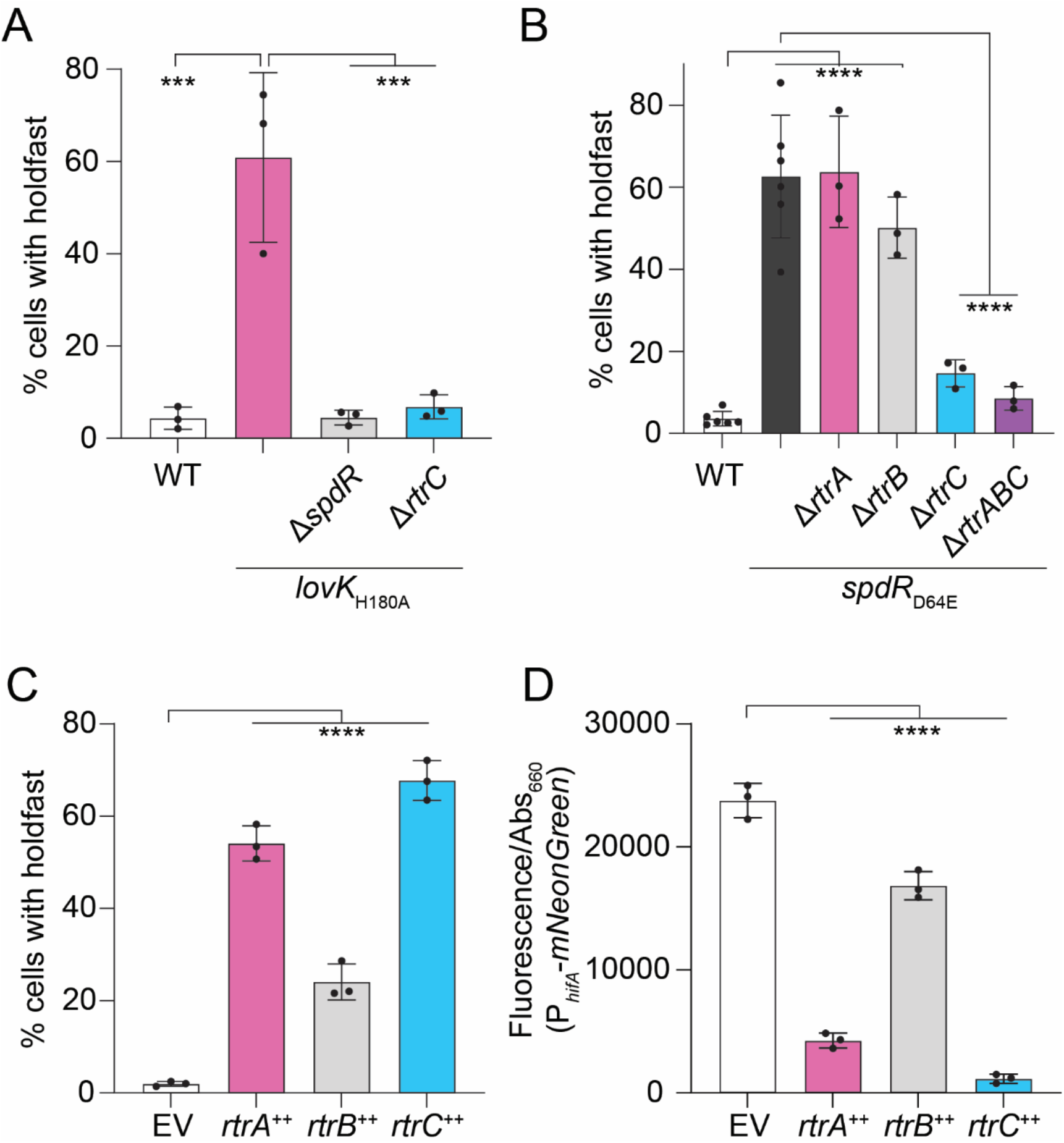
*CCNA_00551* (*rtrC*) regulates holdfast synthesis and *hfiA* expression. **A)** Percentage of cells with stained holdfast in wild type (WT) or *lovK*_H180A_ strains bearing in-frame deletions (Δ) in *spdR* and *CCNA_00551* (*rtrC*). **B)** Percentage of cells with stained holdfast in WT, *spdR*_D64E_, or *spdR*_D64E_ strains bearing in-frame deletions in *rtrA, rtrB*, and *rtrC*. **C)** Percentage of cells with stained holdfasts in empty vector (EV), *rtrA, rtrB*, and *rtrC* overexpression (++) backgrounds. **A-C)** Stained holdfasts were quantified by fluorescence microscopy. **D)** *hfiA* transcription in EV, *rtrA, rtrB*, and *rtrC* overexpression (++) backgrounds as measured using a P_*hfiA*_-*mNeonGreen* fluorescent reporter. **A-D)** Strains were grown in M2-xylose defined medium. Fluorescence was normalized to cell density; data show the mean fluorescence. Error bars are standard deviation of three biological replicates, except WT and *spdR*_D64E_ in panel B, which have six biological replicates. Statistical significance was determined by one-way ANOVA followed by **A-B)** Tukey’s multiple comparisons test or **C-D)** Dunnett’s multiple comparison (p-value < 0.001,***; p-value < 0.0001,****).

We expected that performing this genetic selection in a hyper-holdfast *lovK*_H180A_ background would not only uncover previously identified loss-of-adhesion mutants [22] but would also identify new regulators that function to activate holdfast synthesis downstream of LovK. As expected, strains harboring transposon insertions in all the known adhesion TCS genes (e.g. *lovK, spdS, spdR* and *skaH*) had increased abundance in the supernatant (i.e. decreased adhesion to cheesecloth, and positive fitness scores) when grown in the presence of cheesecloth. Insertions in select polar development regulators, and in holdfast synthesis and anchoring genes also resulted in the expected positive fitness scores (Figures 1C and Table S1). Strains with insertions in gene locus *CCNA_00551*, which encodes a predicted standalone 146-residue hypothetical protein, had strongly positive fitness scores after cheesecloth selection. In fact, strains with insertions in *CCNA_00551* were enriched in the supernatant to a greater extent than TCS adhesion mutants or *rtrA* and *rtrB* mutants (Figure 1C; Table S1). Consistent with these Tn-seq data, in-frame deletion of either *spdR* or *CCNA_00551* from the chromosome abrogated the hyper-holdfast phenotype of *lovK*_H180A_ (Figure 2A). Expression of *CCNA_00551* is directly activated by the DNA-binding response regulator, SpdR [16, 23], which implicated *CCNA_00551* in the adhesion TCS pathway. Following the convention of previously named adhesion factors that function downstream of SpdR [16], we henceforth refer to *CCNA_00551* as *rtrC*.

SpdR functions downstream of LovK [16] (Figure 1A) and expression of a phosphomimetic allele of SpdR (SpdR_D64E_) provides an alternative genetic approach to constitutively activate the *C. crescentus* adhesion TCS system. We predicted that deletion of *rtrC* would also abrogate the hyperadhesive phenotype of a *spdR*_D64E_ strain. Consistent with this prediction and with the Tn-seq data, we observed that the fraction of cells with visibly stained holdfasts was reduced in a *spdR*_D64E_ Δ*rtrC* strain compared to the *spdR*_D64E_ parent (Figure 2B). There was no significant difference in the percentage of cells with visibly stained holdfasts between *spdR*_D64E_ Δ*rtrA* Δ*rtrB* Δ*rtrC* and *spdR*_D64E_ Δ*rtrC* (Figure 2B). This provides evidence that RtrC is the primary downstream determinant of hyperadhesion when the TCS adhesion pathway is constitutively active. Indeed, overexpression of *rtrC* alone enhanced the fraction of cells with stained holdfasts more than overexpression of either *rtrA* or *rtrB* (Figure 2C).

### RtrC is a predicted transcription factor

A search of protein domain family databases in InterPro [24] and the Conserved Domain Database [25] failed to identify conserved domains in RtrC. However, a primary and secondary structure profile matching approach [26] indicated that RtrC resembled classic transcription factors. To explore this possibility, we implemented AlphaFold [27] to predict the tertiary structure of RtrC. This approach predicted a fold that contained five α-helices (α1 – α5) and two β-strands (β1 – β2) that form an antiparallel β hairpin (Figure 3A). We compared this structure to the Protein Data Bank (PDB) using Dali [28], which revealed that the predicted structure of RtrC was most similar to MepR (PDB: 3ECO), a MarR-family transcriptional regulator from *Staphylococcus aureus* containing a winged helix-turn-helix motif [29]. Based on the structural alignments and 3D superposition with MepR, α1 and α5 of RtrC likely form a dimerization domain, while α2, α3, α4, β1, and β2 form a winged helix-turn-helix (Figure 3). Considering these structural predictions, we hypothesized that *rtrC* encoded a transcription factor that functions downstream of the *C. crescentus* TCS adhesion regulatory system.

**Figure 3.**
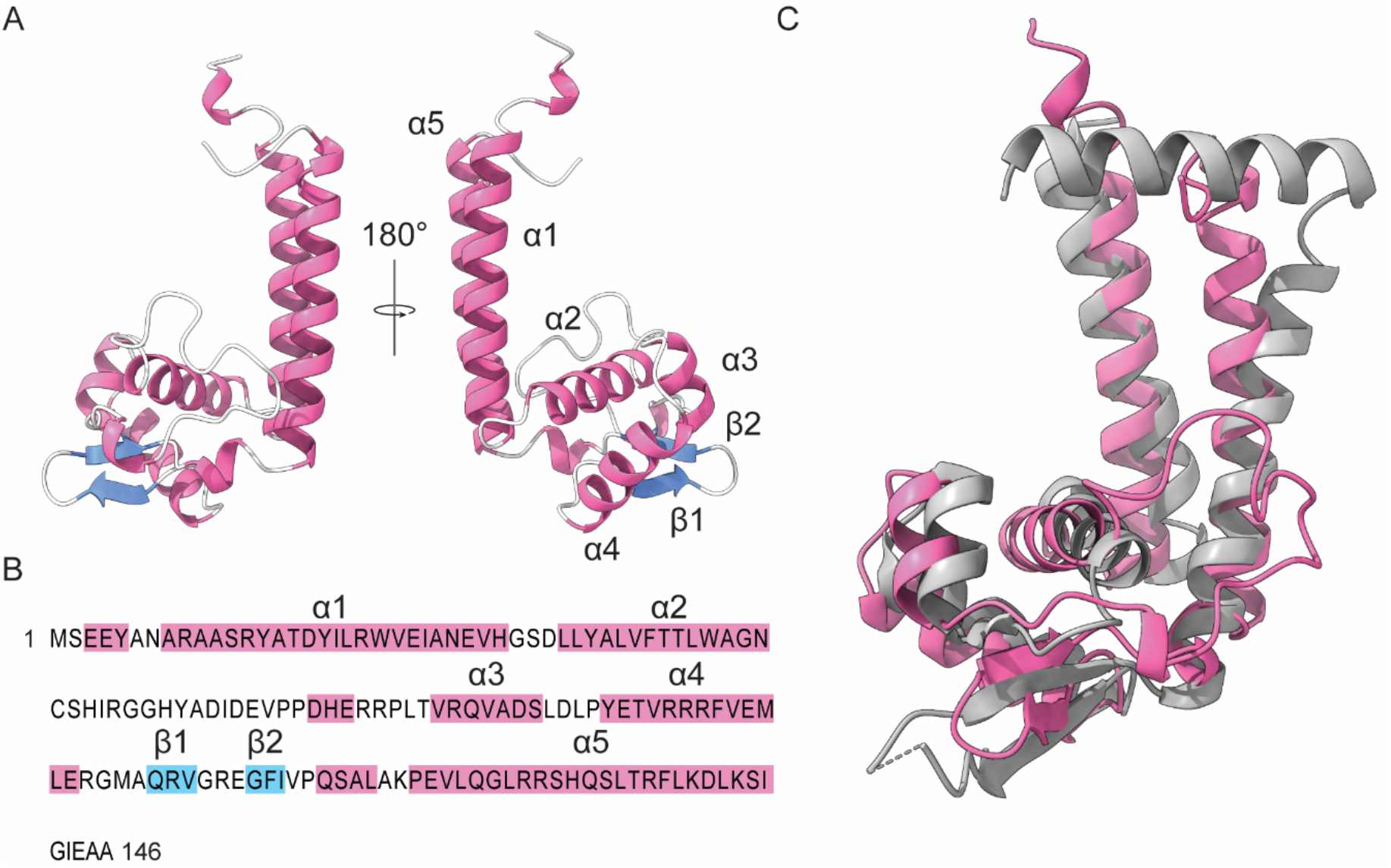
Structural analysis of RtrC. **A)** Tertiary structure of RtrC predicted by AlphaFold [27]. Pink ribbons indicate alpha (α) helices and blue arrows indicate beta (β) strands. Labels (α1-5 and β1-2) correspond to the sequence highlighted in panel B. **B)** Primary structure of RtrC. Amino acids highlighted in pink are in predicted α-helices and residues highlighted in blue are in predicted β-strands, as labeled above the sequence. **C)** RtrC and MepR superposition. RtrC predicted structure is colored pink and MepR (PDB: 3eco – chain A) is colored grey. Dashed lines indicate missing structure. Superposition performed with the Dali server (Z-score: 10.5, rmsd: 3.3) [28].

### RtrC is a potent repressor of the holdfast inhibitor, hfiA

The transcription factors RtrA and RtrB are known to activate holdfast synthesis and adhesion by repressing transcription of the holdfast inhibitor, *hfiA* [16]. Given the correlated phenotypes of *rtrA, rtrB*, and *rtrC* mutants and the prediction that RtrC is a transcription factor (Figures 2 & 3), we hypothesized that RtrC functioned as a transcriptional repressor of *hfiA*. To test this model, we measured changes in expression from a fluorescent *hfiA* transcriptional reporter upon overexpression of *rtrC*. As expected, overexpression of *rtrA* and *rtrB* reduced signal from the P_*hfiA*_ fluorescent reporter by 80% and 30%, respectively. Overexpression of *rtrC* resulted in a 95% reduction in *hfiA* expression (Figure 2D).

### RtrC binds to a pseudo-palindromic DNA motif in vivo and in vitro

We next sought to directly test the predicted DNA-binding function of RtrC. We performed chromatin immunoprecipitation sequencing (ChIP-seq) using a 3xFLAG-tagged *rtrC* allele and identified 113 statistically significant peaks across the genome (Table S2). As expected, we observed significant peak within the *hfiA* promoter region (Figure 4A). Peaks were highly enriched near globally defined transcription start sites (TSS) [30-32] when compared to a set of randomly generated peaks (Figure 4B); this TSS-proximal enrichment pattern is characteristic of proteins that directly bind DNA to regulate gene expression. To identify putative binding motifs in the ChIP-seq peaks, we analyzed the peak sequences using the XSTREME algorithm within the MEME Suite [33]. This revealed a pseudo-palindromic motif in 112 of the 113 *rtrC* peaks (E-value: 2.3e^-12^) that likely corresponded to an RtrC binding site (Figure 4C).

**Figure 4.**
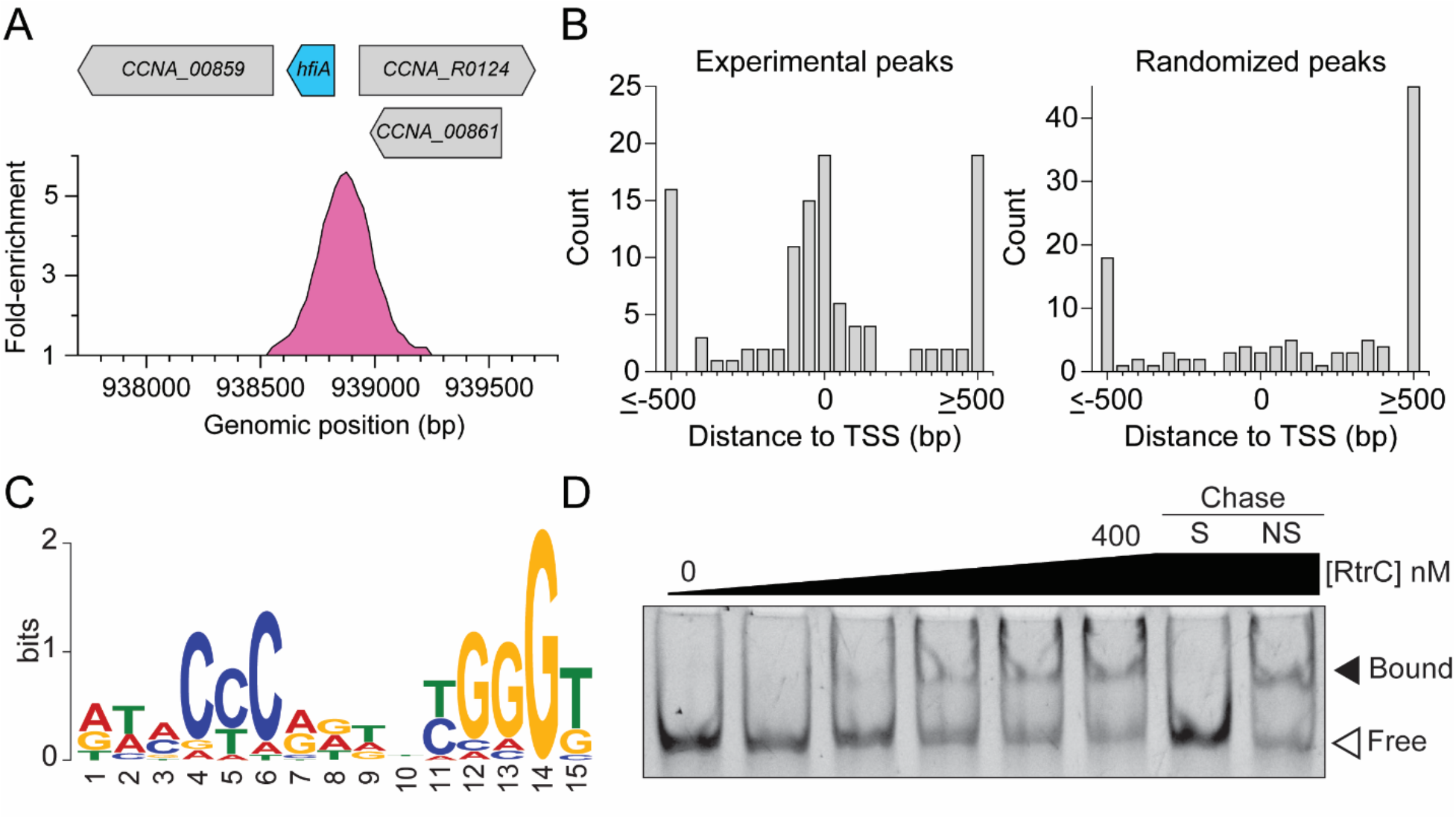
RtrC binds DNA *in vivo* and *in vitro*. **A)** RtrC binds the *hfiA* promoter *in vivo*. ChIP-seq profile from RtrC-3xFLAG pulldowns were plotted as fold-enrichment in read counts compared to the input control. Genomic position of the binding peak (in pink) on the *C. crescentus* chromosome and relative gene locations are marked. **B)** Distribution of RtrC peaks relative to experimentally defined transcription start sites (TSS). Distance from summit of RtrC ChIP-seq peak or randomized peaks to the nearest TSS (113 peaks) were analyzed and are plotted as a histogram. **C)** DNA sequence motif enriched in RtrC ChIP-seq peaks identified by XSTREME [33]. **D)** Electrophoretic mobility shift assay using purified RtrC and *hfiA* promoter sequence. Increasing concentrations of purified RtrC (0, 50, 100, 200, 300, and 400 nM) were incubated with 6.25 nM labeled *hfiA* probe. Specific chase (S) and non-specific chase (NS) contained 2.5 μM unlabeled *hfiA* probe and unlabeled shuffled *hfiA* probe, respectively. Blot is representative of two biological replicates.

To test if RtrC bound to this predicted binding site, we performed electrophoretic mobility shift assays (EMSA) with purified RtrC. Increasing concentrations of RtrC shifted a labeled DNA probe, containing a 27 bp sequence from the *hfiA* promoter centered on the predicted RtrC binding motif (Figure S1 & 4D). RtrC bound to this pseudo-palindrome in the *hfiA* promoter with high affinity (k_d_ of 45 ± 9 nM) (Figure S2). Addition of excess unlabeled specific DNA probe competed with labeled probe bound to RtrC, while unlabeled non-specific probe did not compete for RtrC binding (Figure 4D). These data provide evidence that RtrC directly represses *hfiA* transcription by specifically binding to a pseudo-palindromic motif in the *hfiA* promoter.

### RtrC is a transcriptional activator and repressor

To further characterize the function of RtrC as a transcriptional regulator, we used RNA sequencing (RNA-seq) to measure changes in transcript levels upon *rtrC* overexpression (*rtrC*^++^). RNA-seq was performed with an *rtrC* overexpression strain rather than a *rtrC* deletion strain because *rtrC* expression is low under standard logarithmic growth conditions [30]. By combining RNA-seq and ChIP-seq datasets, we identified genes that are directly controlled by RtrC. Direct targets were defined as genes that *a)* were differentially regulated in *rtrC*^*++*^ relative to an empty vector control, *b)* contained an RtrC-enriched peak by ChIP-seq, and *c)* contained an RtrC binding motif in their promoter region [32]. Of the directly regulated genes, 63% were activated and 37% were repressed by RtrC (Figure 5A; Table S3). Consistent with transcriptional reporter analysis (Figure 2D), *hfiA* transcript levels were ∼5-fold lower in *rtrC*^++^ compared to the vector control (Table S3). To confirm the RNA-seq results, we constructed several fluorescent transcriptional reporters for genes identified as direct targets of RtrC. Consistent with the RNA-seq data, *rtrC* overexpression significantly increased reporter signal for *CCNA_00629* (2.6-fold) and *CCNA_00538* (2.0-fold) and decreased reporter signal for *CCNA_00388* (6.7-fold) compared to an empty vector control (Figure 5B). RtrC bound to the *rtrC* promoter *in vivo* as demonstrated by ChIP-seq, and signal from a *rtrC* transcriptional reporter was 19-fold lower when *rtrC* was overexpressed (Figure 5B). From this, we conclude that RtrC is a negative autoregulator.

**Figure 5.**
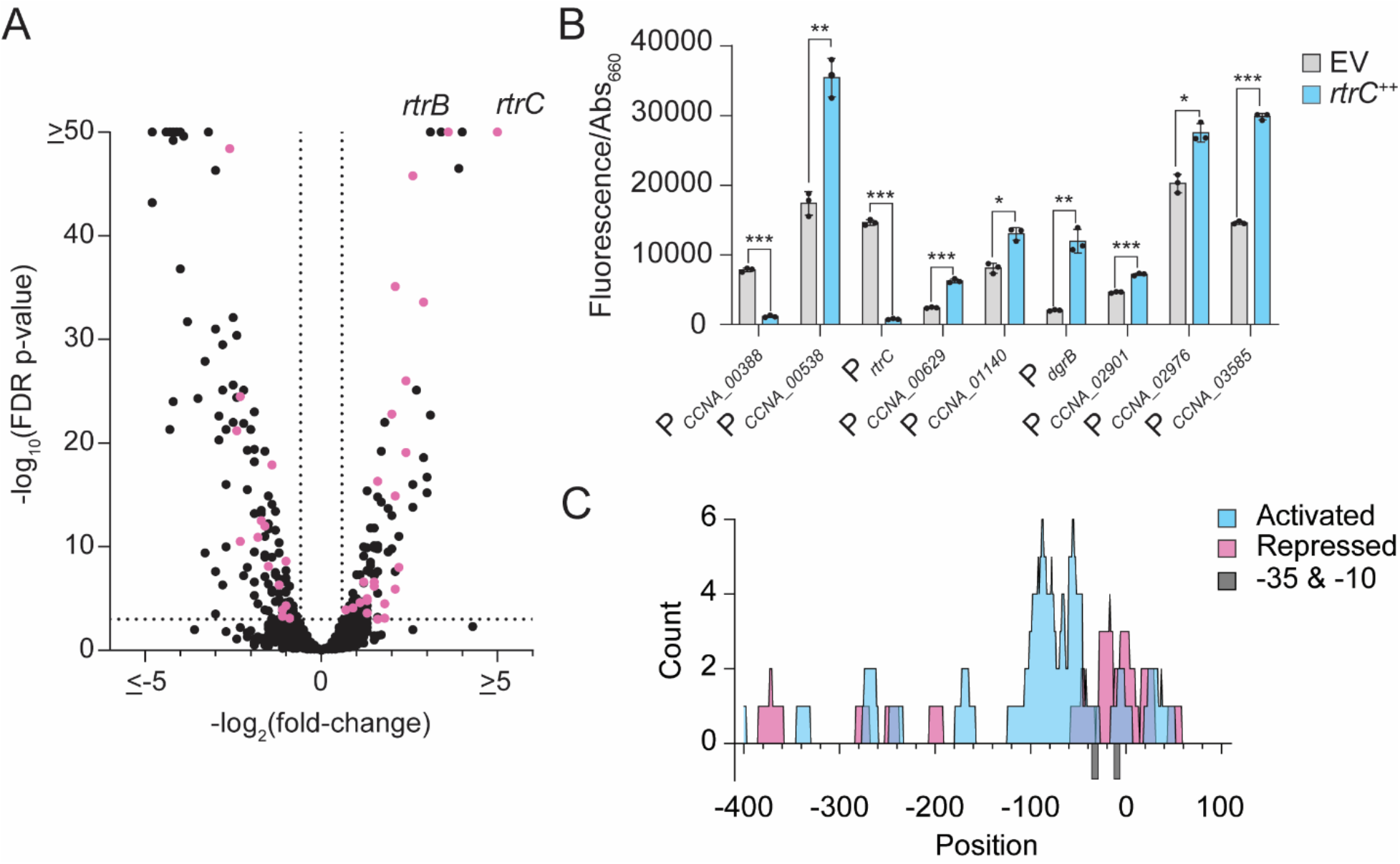
RtrC functions as a transcriptional activator and repressor. **A)** RNA-seq analysis of genes significantly regulated upon *rtrC* overexpression. Volcano plot showing log_2_(fold-change) in transcript levels in an *rtrC* overexpression strain (*rtrC*^++^) versus empty vector (EV) are plotted against -log_10_(FDR p-value). Black dots indicate genes without RtrC motifs and pink dots indicate genes with RtrC motifs in their promoters. Data calculated from four biological replicates. **B)** Transcription from predicted RtrC-regulated promoters measured by promoter fusions to *mNeonGreen*. Cells grown in complex medium (PYE) and fluorescence measured in *rtrC*^++^ or empty vector (EV) backgrounds was normalized to cell density (OD_660_). Data show the mean signal; error bars are standard deviation of three biological replicates. Statistical significance was determined by multiple unpaired t tests, correcting for multiple comparisons using the Holm-Šídák method (p-value < 0.05,*; p-value < 0.01,**; p-value < 0.001,***). **C)** Activity of RtrC as a transcriptional activator or repressor correlates with position of the pseudo-palindromic RtrC motif in the promoter. Distribution of RtrC motifs in promoters (−400 to +100 bp from the transcription start site; TSS) directly regulated by RtrC. Position of RtrC motifs relative to the TSS in each promoter were plotted. Blue indicates motif positions in promoters activated by RtrC (n=26) and pink indicates motif positions in promoters repressed by RtrC (n=16). Grey bars below the x-axis indicate the -35 and -10 positions relative to the annotated TSS.

We also measured signal from transcriptional reporters for several genes that contained RtrC motifs in their promoters but did not meet the statistical threshold for differential regulation by *rtrC* overexpression in the RNA-seq dataset. We evaluated these additional reporters in complex medium to better match conditions in which we identified RtrC-binding peaks. However, for most reporters (11/16) we still observed no significant transcriptional response to *rtrC* overexpression (Figure S3). *rtrC* overexpression significantly enhanced transcription from the remaining 5 reporters: *CCNA_03585* (2.0-fold), *CCNA_02901* (1.6-fold), *dgrB* (5.8-fold), *CCNA_01140* (1.6-fold), and *CCNA_02976* (1.4-fold) (Figure 5B). Together, these ChIP-seq, RNA-seq and reporter data provide evidence that RtrC can function as both a direct transcriptional activator and repressor.

### RtrC motif position within regulated promoters correlates with transcriptional activity

We hypothesized that the activity of RtrC as an activator or repressor depends on its binding position within a promoter relative to the transcription start site (TSS). To assess whether position correlated with regulatory activity, we analyzed the location of RtrC binding motifs within the promoters of genes that were up- or downregulated based on RNA-seq and transcriptional reporter data. Promoters directly repressed by RtrC typically had predicted motifs that overlapped the -10/-35 region of the promoter. In contrast, genes activated by RtrC had binding motifs that were located upstream of the -10/-35 region (Figure 5C). These data provide evidence that the regulatory activity of RtrC is related to the position of the RtrC binding site in a promoter. The results of this analysis are consistent with a well-described trend in which DNA-binding regulators that function as repressors bind at or near the transcription start site, while activators typically bind upstream of the -10/-35 region to promote transcription [34].

### SpdR, RtrB, and RtrC form an OR-gated Type I coherent feedforward loop

Transcript levels of *rtrB* were 12-fold higher in the *rtrC*^++^ background relative to a vector control, placing it among the most highly activated direct targets of RtrC (Table S3). As noted above, SpdR activates transcription of both *rtrB* and *rtrC* [16, 23]. This suggested that these three proteins form a coherent type I feedforward loop (FFL) because the sign of direct regulation (i.e. activation of *rtrB* by SpdR) is the same as the sign of the indirect regulation (i.e. activation of *rtrB* by SpdR through RtrC) (Figure 6A). The regulatory properties of this predicted coherent type I FFL depend on whether *C. crescentus* uses AND-gated logic, in which both SpdR and RtrC are required to activate *rtrB* expression, or OR-gated logic, in which either SpdR or RtrC can activate *rtrB* expression [35]. To test FFL gating, we deleted *spdR* and *rtrC* from the chromosome and measured fluorescence from a *rtrB* transcriptional reporter upon expression of *spdR*_D64E_ and/or *rtrC* from inducible promoters. Expression of either *rtrC* or *spdR*_D64E_ alone increased transcription from the *rtrB* reporter by ∼5-fold, while expression of both *rtrC* and *spdR*_D64E_ increased transcription by ∼6-fold (Figure 6B). *spdR* deletion significantly reduced transcription from a P_*rtrB*_ reporter in stationary phase (Figure S4). As expected, deletion of *rtrC* alone did not affect transcription from P_*rtrB*_ as *spdR* is still present on the chromosome (Figure S4). We conclude that either SpdR or RtrC can activate *rtrB* expression and are therefore competent to form an OR-gated coherent type I FFL in *C. crescentus*.

**Figure 6.**
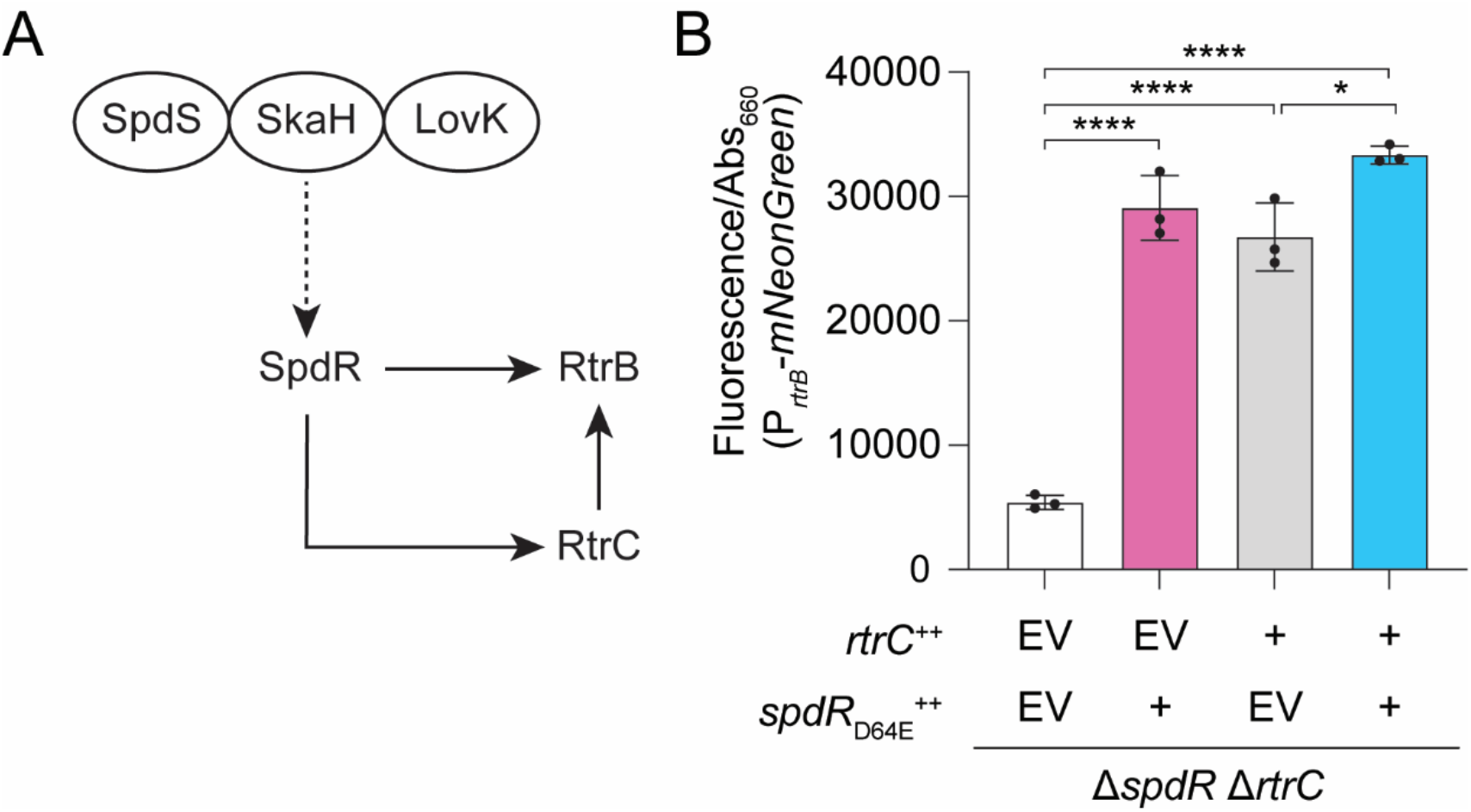
*spdR*-*rtrC*-*rtrB* form an OR-gated type I coherent feedforward loop. **A)** Schematic of the Type I coherent feedforward loop. The sensor histidine kinases SpdS, SkaH, and LovK function upstream and regulate the DNA-binding response regulator, SpdR [16]. SpdR can activate transcription of both *rtrC* and *rtrB*; RtrC activates transcription of *rtrB*. Dashed arrows indicate post-transcriptional activation and solid arrows indicate transcriptional activation. **B)** *rtrB* transcription measured using a P_*rtrB*_-*mNeonGreen* transcriptional reporter. Reporter strains were built in a background in which *rtrC* and *spdR* are deleted from the chromosome (Δ*spdR* Δ*rtrC*). Transcription was measured in empty vector (EV), *rtrC*, or *spdR*_D64E_ overexpression (++) backgrounds. Cells were grown in complex medium (PYE) and fluorescence signal was normalized to cell density (OD_660_). Data are the mean; errors bars represent standard deviation of three biological replicates. Statistical significance was determined by one-way ANOVA followed by Tukey’s multiple comparisons test (p-value < 0.05,*; p-value < 0.0001,****).

### rtrC mutants have altered adhesion profiles in cellulosic substrate binding and pellicle assays

SpdR affects gene regulation during stationary phase [23, 36], and was previously reported to bind *rtrC* promoter DNA [23]. Consistent with these observations, transcription from a *rtrC* reporter increased 13-fold during stationary phase in complex medium in a *spdR*-dependent manner (Figure S5). The regulation of *rtrC* transcription is strongly medium dependent as stationary phase activation of *rtrC* was not observed in M2-xylose defined medium (Figure S5). These results led us to assess the effect of *rtrC* gene deletion on holdfast synthesis in log and stationary phase cultures in complex medium. The fraction of cells with holdfasts in complex medium during early log phase was not significantly different in strains with in-frame deletions of *spdR, rtrA, rtrB, rtrC*, or in a strain missing all three *rtr* regulators (Figure S6A). Stained holdfast were greatly reduced in stationary phase, but this effect did not require *spdR*. Holdfast counts are low in stationary phase, and an *rtrB* deletion mutant had even fewer holdfasts than wild type in stationary phase; deletion of either *rtrA* or *rtrC* had no effect on holdfast counts under these conditions (Figure S6B). While Δ*rtrC* holdfast counts were not significantly different from wild type in standard complex medium cultures, analysis of transposon sequencing data revealed that strains harboring insertions in *rtrC* are highly enriched in the supernatant after repeated passaging in media containing cheesecloth (Figure 7A, data from [22]). Thus, there is evidence that loss of *rtrC* function results in diminished adherence to a solid cellulosic substrate after repeated passage. Importantly, disruption of other genes in the TCS adhesion regulation pathway resulted in a similar temporal adhesion profile as *rtrC::Tn* mutants in this serial passage experiment (Figure 7A & 7D; Figure S7A).

**Figure 7.**
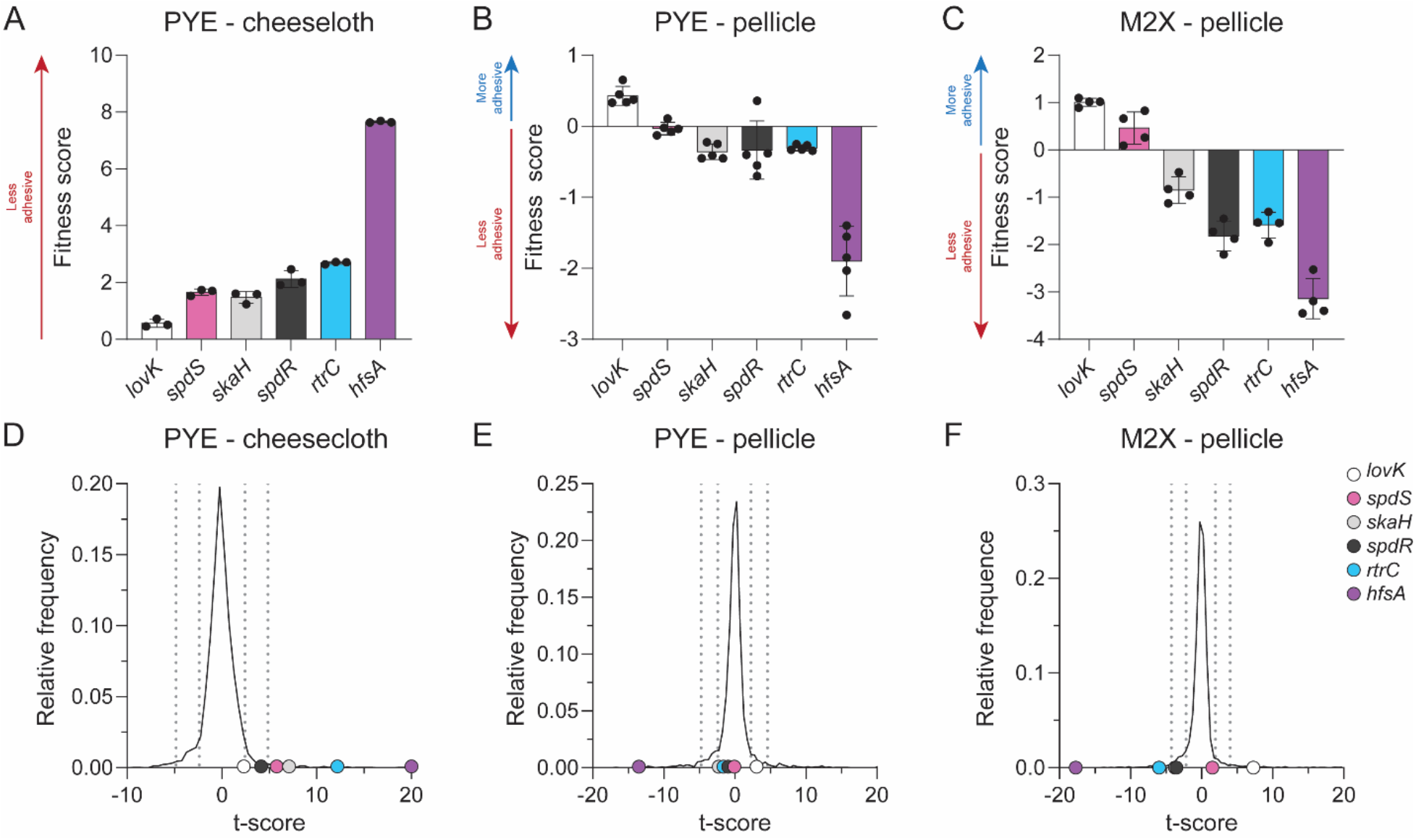
Mutant fitness profiles in cheesecloth adherence and pellicle assays. **A-C)** Composite fitness scores of *C. crescentus* strains harboring transposon insertions in adhesion regulatory genes (*lovK, spdS, skaH, spdR*, and *rtrC*) and a representative holdfast synthesis gene (*hfsA*) after **(A)** 5 days of serial passaging in the presence of cheesecloth where strains were sampled from the supernatant, which is enriched with non-adherent cells [22], (**B)** 4 days of static cultivation in complex (PYE) medium where strains were sampled from the pellicle at the air-liquid interface, or (**C)** 7 days of static cultivation in M2-xylose defined medium where strains were sampled from the pellicle at the air-liquid interface. Data in panels A-C show the mean ± standard deviation, with experimental replicates plotted as points (n=3 to 5 depending on the experiment). **D-F)** Frequency distribution of mean t-value for each gene in the full dataset after **(D)** passaging in the presence of cheesecloth, **(E)** static growth in complex (PYE) medium, or **(F)** static growth in M2-xylose defined medium. The t-value is the fitness value of a gene divided by a variance metric based on the total number of reads for each gene (as previously described [61]) and provides a metric to assess the significance of mutant fitness values. Labeled dots mark the t-values of adhesion regulator mutants (*lovK, spdS, skaH, spdR*, and *rtrC*) and a representative holdfast synthesis gene (*hfsA*). Dotted vertical lines mark boundaries that contain 68%, and 95% of the t-values in each experiment, which would reflect one and two standard deviations in a normally distributed set.

In addition to its critical role in adhesion to solid surfaces, the holdfast is also required for *C. crescentus* to form pellicle biofilms at air-liquid interfaces [37]. We therefore hypothesized that strains with disruptions in genes that can promote holdfast formation, including *rtrC*, would have reduced abundance in the pellicle micro-environment. To test this model, we grew the same barcoded transposon mutant library used in the cheesecloth adhesion experiments [22, 38] in either defined or complex medium under static growth conditions, which promotes pellicle formation. *C. crescentus* strains were sampled from the air-liquid interface at successive stages of pellicle development. Strain barcodes were then amplified, sequenced, counted, and fitness scores were calculated for each gene. Positive fitness scores indicate mutant strain enrichment in the pellicle fraction and negative scores reflect underrepresentation of mutants in the pellicle. As expected, strains harboring disruptions of genes required for holdfast synthesis (e.g. *hfsA*) were highly underrepresented in pellicles in both defined and complex media (Figure 7B-C & 7E-F & S7B-C; Table S4). Transposon insertions in *rtrC* resulted in only a minor reduction in strain abundance in the pellicle in complex medium, similar to strains with insertions in the TCS adhesion regulators *skaH* and *spdR*. Strains harboring insertions in *lovK* were slightly enriched in the pellicle (Figure 7B & 7E). We conclude that in complex medium, the adhesion signaling pathway only weakly contributes to holdfast-dependent pellicle formation. However, in M2-xylose defined medium *rtrC*::*Tn* mutants have reduced abundance in pellicles, again similar to *skaH* and *spdR* mutants (Figure 7C & 7F), and the effect of *lovK* disruption is more pronounced and is consistent with *lovK* playing a repressive role in adhesion TCS signaling under this condition. This result echoes the repressive effect that *lovK* has in regulation of the general stress response (GSR) [39], and may be related to our observation that disruption of the core GSR regulators, *phyR* and *ecfG*, attenuates the hyperadhesive phenotype of *lovK*_H180A_ (Figure 1C). Taken together, these results are consistent with a model in which the adhesion TCS regulatory system is active in static growth in defined xylose medium. RtrC, a downstream component of the TCS adhesion pathway that directly represses *hfiA*, plays a role pellicle development in defined medium.

## Discussion

We designed a forward genetic selection to search for novel holdfast regulators and identified RtrC. This formerly hypothetical protein functions downstream of an ensemble of TCS regulatory genes to activate surface adhesion. RtrC binds and regulates multiple sites on the *C. crescentus* chromosome, including the *hfiA* promoter where it represses *hfiA* transcription and thereby activates holdfast synthesis.

### RtrC structure and regulatory activity

A comparison of the predicted three-dimensional structure of RtrC to experimental structures available in the PDB suggested structural similarity to MepR and several other MarR family transcriptional regulators (Figure 3C). Members of this transcription factor family often bind as dimers to pseudo-palindromic DNA sequences [40-42]. MarR family transcriptional regulators are known to function as both activators and repressors, depending on the position of binding within regulated promoters. Similarly, we observed that the activity of RtrC as an activator or repressor was correlated with the position of the RtrC motif within the promoter; this positional effect on transcriptional regulation is a well-described phenomenon [43]. Our data thus provide evidence that RtrC (like MarR) functions as a classic transcription factor. The sequence of RtrC is not broadly distributed; it is largely restricted to the Caulobacterales and Rhodospirillales where it is annotated as a hypothetical protein. The genomic neighborhood surrounding *rtrC* is highly conserved across diverse *Caulobacter* species (Figure S8) suggesting *rtrC* is ancestral in the genus.

### A new layer of hfiA regulation

Holdfast-dependent surface attachment in *C. crescentus* is permanent and therefore highly regulated. The small protein, HfiA, is central to holdfast control. It represses holdfast biogenesis by directly interacting with the glycosyltransferase HfsJ, an enzyme required for synthesis of holdfast polysaccharide [15]. *hfiA* expression is influenced by multiple cell cycle regulators, TCS sensory/signaling systems, a transcriptional regulator of stalk biogenesis, and c-di-GMP [15-17, 44]. We have shown that RtrC functions immediately downstream of the stationary phase response regulator, SpdR, to directly bind the *hfiA* promoter and repress its transcription. SpdR can therefore regulate expression of at least three distinct direct repressors of *hfiA* transcription – *rtrA, rtrB*, and *rtrC* (Figure 8).

**Figure 8.**
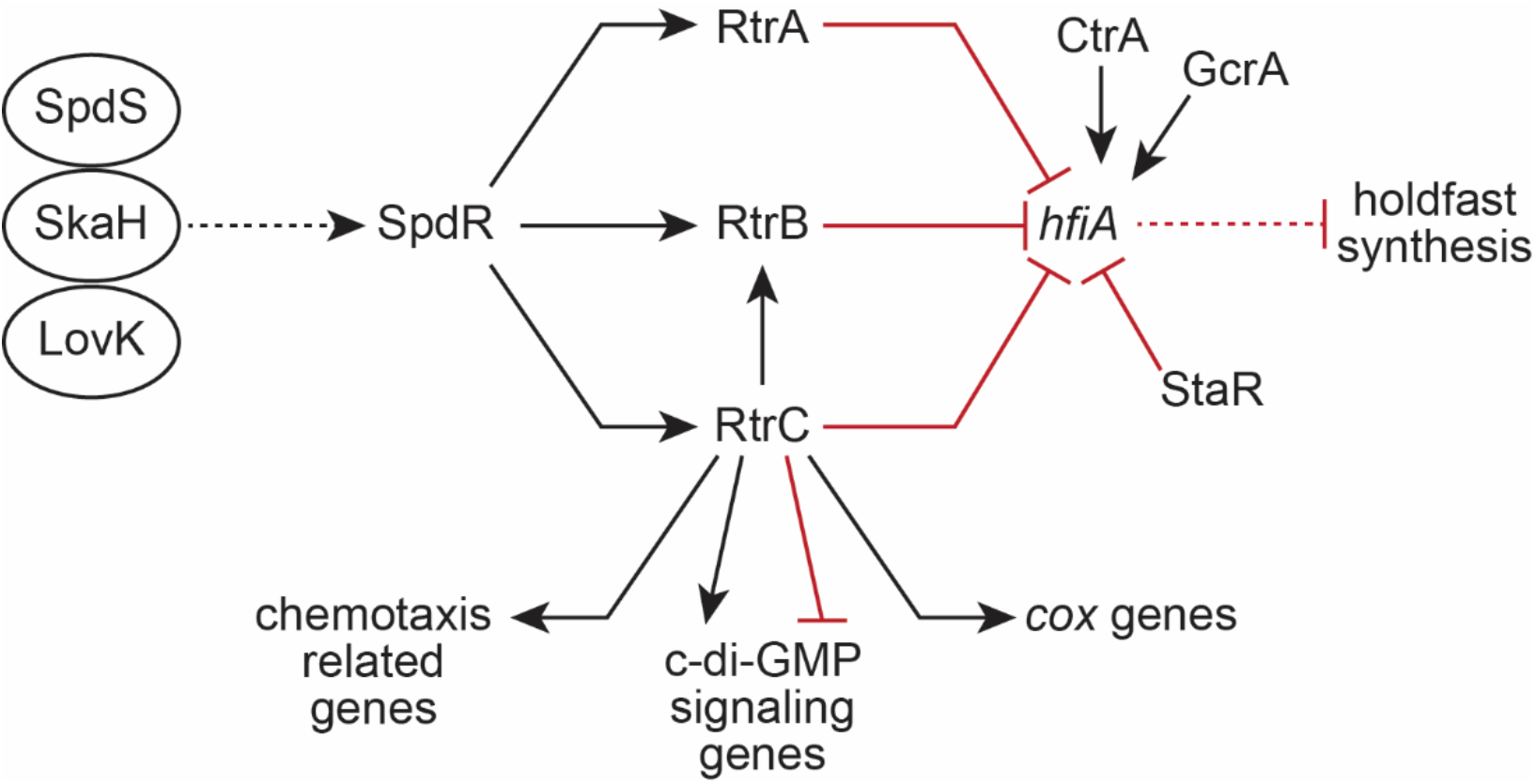
*C. crescentus* adhesion TCS regulatory network. The LovK, SkaH, and SpdS sensor histidine kinases function upstream of the DNA-binding response regulator, SpdR [16]. SpdR directly activates transcription of *rtrA, rtrB*, and *rtrC*. RtrA, RtrB, RtrC, and the XRE-family transcription factor, StaR, all directly repress *hfiA* transcription, while the cell cycle regulators CtrA and GcrA directly activate *hfiA* transcription. In addition to regulating *hfiA*, RtrC can function as a transcriptional activator and repressor for several groups of genes, including those with predicted roles in chemotaxis, c-di-GMP signaling, and aerobic respiration (*cox*). Dashed lines indicate post-transcriptional regulation, solid black arrows indicate transcriptional activation, and red bar-ended lines indicate transcriptional repression.

Why, then, does the *spdR* response regulator have so many outlets to directly modulate *hfiA* transcription? We do not know whether the activities of RtrA, RtrB, or RtrC as transcription factors are allosterically regulated by small molecules, chemical modifications, or protein-protein interactions. If these transcription factors are subject to allosteric regulation, it may be the case that this suite of proteins serves to integrate multiple environmental or cellular signals. In such a model, primary signals that regulate the transcriptional activity of SpdR may enhance expression of RtrA, RtrB, or RtrC, which could then influence the scale of adhesion to substrates in response to secondary physical or chemical cues. Another possible explanation for multiple adhesion transcription factors downstream of SpdR is redundancy. Transcription factor redundancy may buffer the network against transient changes in signaling and gene regulation, ensuring that the decision to synthesize a holdfast (or not) is less subject to environmental fluctuations.

### More on the RtrC regulon

RtrC-dependent regulation of transcription was observed for dozens of genes, suggesting that RtrC influences physiological processes beyond holdfast development. For instance, the *spdR*-*rtrC* axis activates transcription of the *cox* genes (Table S3). These genes encode an aa_3_-type cytochrome oxidase, which is one of four distinct aerobic terminal oxidase complexes in *C. crescentus* [45]. An aa_3_-type oxidase in *Pseudomonas aeruginosa* is reported to provide a survival advantage for cells under starvation conditions [46]. The physiological impact of *cox* regulation by RtrC remains untested.

RtrC directly regulates expression of several genes involved in c-di-GMP (CdG) signaling including a CdG receptor (*dgrB*), a PAS-containing EAL phosphodiesterase (*CCNA_01140*), and a GGDEF-EAL protein (*CCNA_00089*). Deletion of *CCNA_00089* enhances *C. crescentus* surface attachment [47] and *dgrB* is reported to directly bind CdG and repress motility [48]. Considering that *rtrC* overexpression represses *CCNA_00089* expression (Table S3) and activates *dgrB* (Table S3; Figure 5B), it is possible that *rtrC* influences adhesion and/or motility through *CCNA_00089, dgrB* and *hfiA*. RtrC also activates expression of genes with predicted roles in chemotaxis, including two methyl-accepting chemotaxis proteins (*CCNA_00538* and *CCNA_02901*) and a *cheY* receiver domain protein (*CCNA_03585*). Additionally, RtrC activates transcription of an alternative chemotaxis cluster (*CCNA_00628* and *CCNA_00629*-*CCNA_00634*), which has been reported to influence *hfiA* transcription and *C. crescentus* surface adherence [49]. Given these results, it seems likely that *rtrC* will influence *C. crescentus* motility and/or chemotaxis under certain conditions.

We identified RtrC motifs in the promoters of 61 operons that were not differentially regulated in our RNA-seq data. It may be the case that RtrC binding has no effect on the regulation of gene expression at certain sites, or that regulation from particular sites requires additional factors that were not present or expressed under the conditions we assayed. Our data provide evidence that RtrC-dependent gene expression can change as a function of growth medium: *CCNA_00629, CCNA_00538*, and *CCNA_00388* were regulated by RtrC in both defined medium and complex medium as shown by our RNA-seq data and confirmed by transcriptional reporter analysis (Table S3; Figure 5B). In contrast, *CCNA_03585, CCNA_02901, dgrB, CCNA_01140*, and *CCNA_02976* were only regulated by RtrC in complex medium (Figure 5B), while *CCNA_00089* was only regulated in defined medium (Table S3; Figure S3). We do not understand the mechanism(s) underlying media-dependent regulation of components of the RtrC regulon. Published transcriptomic data provide evidence that carbon limitation [50], cell cycle [51], and stringent response signaling [52] all significantly affect *rtrC* transcription, indicating that there are a range of environmental conditions (and developmental states) in which *rtrC* could impact gene expression.

### Signals and feedback control in the adhesion pathway

The DNA-binding response regulator, SpdR, is regulated in a growth phase and media-dependent fashion [23, 36], and systems homologous to *C. crescentus* SpdS-SpdR are reported to respond to cellular redox state and to flux through the electron transport chain via modulation of disulfide bond formation [53], modification of a reactive cysteine [54], or by binding of oxidized quinones [55, 56]. *C. crescentus* SpdS contains both the reactive cysteine and quinone-interacting residues observed in related bacteria, suggesting that SpdS may be regulated in a similar manner. The activity of SpdR as a transcriptional regulator is also affected by the sensor kinases LovK and SkaH [16]. Thus, multiple environmental signals apparently feed into SpdR-dependent gene regulation.

Our data provide evidence that *spdR* and *rtrC* form a type I coherent feedforward loop (C1-FFL) with the XRE-family transcription factor *rtrB*. Experimental and theoretical studies of C1-FFLs indicate that these regulatory motifs function as sign-sensitive delay elements [35, 57]. AND-gated C1-FFLs exhibit a delay in the ON step of output expression, which can allow circuits to function as persistence detectors [35, 58]. Conversely, OR-gated C1-FFLs delay the OFF step of output expression, which can buffer the circuit against the transient loss of activating signals [35, 57]. Expression of either *spdR* or *rtrC* was sufficient to activate transcription from a P_*rtrB*_ reporter, indicating that the *spdR*-*rtrC*-*rtrB* C1-FFL is competent to function as an OR-gated system (Figure 6). Though the exact environmental signals that regulate the adhesion TCS pathway remain undefined, the architecture of the SpdR-RtrC-RtrB circuit suggests that RtrC can reinforce *rtrB* expression in particular environments where the levels of activating signals for SpdR are fluctuating or noisy.

### The contribution of rtrC to complex adhesion phenotypes

It is clear that RtrC expression is impacted by multiple environmental cues. Tn-seq studies show that *rtrC* mutants are enriched in the supernatant of complex medium after serial passaging in the presence of cheesecloth (Figure 7A & S7A) [22]. *C. crescentus* was repeatedly cycled between different growth states over the course of this five day experiment (i.e. stationary to logarithmic phase) as cells were diluted into fresh media each day. Considering that SpdR strongly activates RtrC expression during stationary phase (Figure S5), it seems likely that the adhesion profile of *rtrC* mutants is influenced by growth phase-dependent changes in *rtrC* expression. Importantly, disruptions in the adhesion TCS system (specifically *spdS, spdR*, and *skaH*) resulted in similar temporal enrichment profiles in the supernatant. This provides evidence that the multi-protein signaling system functioning upstream of *rtrC* similarly influences adhesion under this serial passage condition.

The pellicle provides an interesting and ecologically relevant state to further assess the function of *rtrC* and its upstream regulators. In *C. crescentus*, pellicle formation at the air-liquid interface requires holdfast production [37]. Hyperadhesive mutants have accelerated pellicle development and holdfast null mutants are unable to stably partition to this microenvironment [37]. As expected, we observed that disruption of holdfast synthesis genes led to highly reduced fitness within the pellicle fraction (Figure 7B-C & S7B-C; Table S4). In addition, our data provide evidence that disruption of *rtrC* reduces the ability of *C. crescentus* to inhabit this micro-environment. We hypothesize that *rtrC* mutants have reduced fitness in the pellicle fraction because of differences in holdfast development; this is based on our observations that RtrC regulates holdfast formation through *hfiA*. However, it is possible that reduced fitness of *rtrC* mutants in the pellicle fraction is due to other changes in metabolic state or growth rate affected by RtrC.

Interestingly, the impact of *rtrC* disruption on strain fitness in the pellicle was more significant in defined medium than complex medium (Figure 7B-C; Figure 7E-F; Table S4). We observed similar pellicle fitness trends for *lovK, spdR* and *skaH* mutants in defined versus complex media, providing evidence that the adhesion TCS pathway upstream of *rtrC* plays a larger regulatory role in the pellicle in defined medium than in complex medium. Though *rtrC* expression was not activated by *spdR* in defined medium in a standard continuously shaken culture (Figure S5), our results suggest that SpdR and the adhesion TCS pathway is active in statically grown pellicles in defined medium.

This study expands our understanding of a transcriptional network functioning downstream of a suite of TCS proteins that affect surface adherence in *Caulobacter*. The DNA-binding response regulator SpdR regulates expression of at least three transcription factor genes (*rtrA, rtrB*, and *rtrC*) that directly repress the holdfast inhibitor, *hfiA*. Of these three transcription factors, RtrC is the most potent repressor of *hfiA*. However, it is clear from the ChIP-seq and transcriptomic data presented in this study that the regulatory function of RtrC likely extends well beyond *hfiA* and holdfast synthesis (Figure 8). Efforts focused on deciphering the regulatory cues that impact signaling through the adhesion TCS system, and comparative analyses of the SpdR, RtrA, RtrB, and RtrC regulons will provide a more complete understanding of the regulatory logic that underpins the highly complex process of holdfast adhesin development, surface adherence, and pellicle formation in *Caulobacter*.

## Materials and Methods

### Strain growth conditions

*Escherichia coli* was grown in Lysogeny broth (LB) or LB agar (1.5% w/v) at 37°C [59]. Medium was supplemented with the following antibiotics when necessary: kanamycin 50 μg ml^-1^, chloramphenicol 20 μg ml^-1^, oxytetracycline 12 μg ml^-1^, and carbenicillin 100 μg ml^-1^.

*Caulobacter crescentus* was grown in peptone-yeast extract (PYE) broth (0.2% (w/v) peptone, 0.1% (w/v) yeast extract, 1 mM MgSO_4_, 0.5 mM CaCl_2_), PYE agar (1.5% w/v), or M2 defined medium supplemented with xylose (0.15% w/v) as the carbon source (M2X) [60] at 30°C. Solid medium was supplemented with the following antibiotics where necessary: kanamycin 25 μg ml^-1^, chloramphenicol 1 μg ml^-1^, and oxytetracycline 2 μg ml^-1^. Liquid medium was supplemented with the following antibiotics where necessary: chloramphenicol 1 μg ml^-1^, and oxytetracycline 2 μg ml^-1^.

### Tn-Himar mutant library construction and mapping

Construction and mapping of the barcoded Tn-himar library was performed following protocols originally described by Wetmore and colleagues [61]. A 25 ml culture of the *E. coli* APA_752 barcoded transposon donor pool (obtained from Adam Deutschbauer Lab) was grown to mid-log phase in LB broth supplemented with kanamycin and 300 μM diaminopimelic acid (DAP). A second 25 ml culture of *C. crescentus lovK*_H180A_ was grown to mid-log phase in PYE. Cells from both cultures were harvested by centrifugation, washed twice with PYE containing 300 μM DAP, mixed and spotted together for conjugation on a PYE agar plate containing 300 μM DAP. After incubating the plate overnight at room temperature, the cells were scraped from the plate, resuspended in PYE medium, spread onto 20, 150 mm PYE agar plates containing kanamycin and incubated at 30°C for three days. Colonies from each plate were scraped into PYE medium and used to inoculate a 25 ml PYE culture containing 5 μg ml^-1^ kanamycin. The culture was grown for three doublings, glycerol was added to 20% final concentration, and 1 ml aliquots were frozen at -80°C.

To map the sites of transposon insertion, we again followed the protocols of Wetmore et al. [61]. Briefly, genomic DNA was purified from three 1 ml aliquots of each library. The DNA was sheared and ∼300 bp fragments were selected before end repair. A Y-adapter (Mod2_TS_Univ, Mod2_TruSeq) was ligated and used as a template for transposon junction amplification with the primers Nspacer_BarSeq_pHIMAR and either P7_mod_TS_index1 or P7_mod_TS_index2. 150-bp single end reads were collected on an Illumina HiSeq 2500 in rapid run mode, and the genomic insertion positions were mapped and correlated to a unique barcode using BLAT [62] and MapTnSeq.pl to generate a mapping file with DesignRandomPool.pl. Using this protocol, we identified 232903 unique barcoded insertions at 60940 different locations on the chromosome. The median number of barcoded strains per protein-encoding gene (that tolerated Tn insertion) was 34; the mean was 49.6. Median number of sequencing reads per hit protein-encoding gene was 4064; mean was 6183.5. All code used for this mapping and analysis is available at https://bitbucket.org/berkeleylab/feba/.

### Adhesion profiling of the lovK_H180A_ Tn-Himar mutant library

Adhesion profiling followed the protocol originally outlined in Hershey et al. [22]. 1 ml aliquots of the barcoded transposon library were cultured, collected by centrifugation, and resuspended in 1 ml of M2X medium. 300 μl of this barcoded mutant pool was inoculated into a well of a 12-well microtiter plate containing 1.5 ml M2X defined medium with 6-8 ∼1 × 1 cm layers of cheesecloth. These microtiter plates were incubated for 24 hours at 30°C with shaking at 155 rpm after which 150 μl of the culture was passaged by inoculating into a well with 1.65 ml fresh M2X containing cheesecloth. Cells from an additional 500 μl of medium from each well was harvested by centrifugation and stored at -20°C for barcode sequencing (BarSeq) analysis. Each passaging experiment was performed in triplicate, and passaging was performed sequentially for a total of five rounds of selection. Identical cultures grown in a plate without cheesecloth were used as a nonselective reference condition.

Cell pellets were used as PCR templates to amplify the barcodes in each sample using indexed primers [61]. Amplified products were purified and pooled for multiplexed sequencing. 50 bp single end reads were collected on an Illumina HiSeq4000. The Perl and R scripts MultiCodes.pl, combineBarSeq.pl and FEBA.R were used to determine fitness scores for each gene by comparing the log_2_ ratios of barcode counts in each sample over the counts from a nonselective growth in M2X without cheesecloth. To evaluate mutant phenotypes in each selection, the replicates were used to calculate a mean fitness score for each gene after each passage. Mean fitness (a proxy for adhesion to cheesecloth) was assessed across passages for each gene.

### Plasmid and strain construction

Plasmids were cloned using standard molecular biology techniques and the primers listed in Table S5. To construct pPTM051, *CCNA_03380* (−21 to +15 bp relative to the start of the gene) was fused to *mNeonGreen* and cloned into pMT805 lacking the xylose-inducible promoter [63]. To construct pPTM056, site directed mutagenesis was used to introduce a silent mutation in the chloramphenicol acetyltransferase gene of pPTM051 to remove an EcoRI site. A cumate-inducible, integrating plasmid was constructed by fusing a backbone with a chloramphenicol resistance marker derived from pMT681 [63], the xylose integration site derived from pMT595 [63], and the cumate-responsive repressor and promoter derived from pQF through Gibson Assembly [64]. To construct pPTM057, the xylose integration site, cumate repressor, and cumate-inducible promoter of pPTM052 were fused to a backbone with a kanamycin resistance marker derived from pMT426 [63]. For reporter plasmids, inserts were cloned into the replicating plasmid pPTM056. For overexpression constructs, inserts were cloned into pPTM057 or pMT604 that integrate at the xylose locus and contain either a cumate-(P_Q5_) or xylose-inducible (P_xyl_) promoter, respectively [63]. For 3xFLAG-tagged RtrC overexpression, inserts were cloned into the replicating plasmid pQF [64]. Deletion inserts were constructed by overlap PCR with regions up- and downstream of the target gene and cloned into the pNPTS138 plasmid. Clones were confirmed with Sanger sequencing.

Plasmids were transformed into *C. crescentus* by either electroporation or triparental mating [60]. Transformants generated by electroporation were selected on PYE agar supplemented with the appropriate antibiotic. Strains constructed by triparental mating were selected on PYE agar supplemented with the appropriate antibiotic and nalidixic acid to counterselect against *E. coli*. Gene deletions and allele replacements were constructed using a standard two-step recombination/counter-selection method, using *sacB* as the counterselection marker. Briefly, pNPTS138-derived plasmids were transformed into *C. crescentus* and primary integrants were selected on PYE/kanamycin plates. Primary integrants were incubated overnight in PYE broth without selection. Cultures were plated on PYE agar plates supplemented with 3% (w/v) sucrose to select for recombinants that had lost the plasmid. Mutants were confirmed by PCR amplification of the gene of interest from sucrose resistant, kanamycin sensitive clones.

### Holdfast imaging and quantification

Strains were inoculated in triplicate in M2X or PYE, containing 50 μM cumate when appropriate, and grown overnight at 30°C. Strains were subcultured in M2X or PYE, containing 50 μM cumate when appropriate, and grown for 3-8 hours at 30°C. Cultures were diluted to 0.0000057 – 0.00045 OD_660_ and incubated at 30°C until reaching 0.05 – 0.1 OD_660_,. For stationary phase cells, cultures were diluted to 0.05 OD_660_ and incubated at 30°C for 24 hours. Alexa594-conjugated wheat germ agglutinin (WGA) (ThermoFisher) was added to the cultures with a final concentration of 2.5 μg ml^-1^. Cultures were shaken at 30°C for 10 min at 200 rpm. Then, 1.5 ml early log phase culture or 0.75 ml stationary phase culture was centrifuged at 12,000 x g for 2 min and supernatant was removed. Pellets from early log phase in M2X and PYE were resuspended in 35 μl M2X or 100 μl H_2_O, respectively. Pellets from stationary phase in PYE were resuspended in 400 μl H_2_O. Cells were spotted on 1% (w/v) agarose pads in H_2_O and imaged with a Leica DMI6000 B microscope. WGA staining was visualized with Leica TXR ET (No. 11504207, EX: 540-580, DC: 595, EM: 607-683) filter. Cells with and without holdfasts were enumerated using the image analysis suite, FIJI. Statistical analysis was carried out in GraphPad 9.3.1.

### Structure prediction and comparison

The structure of CCNA_00551 was predicted with AlphaFold [27] through Google Colab using the ChimeraX interface [65]. The predicted structure from AlphaFold was submitted to the Dali server [28] for structural comparison to the Protein Data Bank (PDB).

### Chromatin immunoprecipitation sequencing (ChIP-seq)

Strains were incubated in triplicate at 30°C overnight in 10 ml PYE supplemented with 2 μg/ml oxytetracycline. Then, 5 ml overnight culture was diluted into 46 ml PYE supplemented with 2 μg/ml oxytetracycline, grown at 30°C for 2 hours. Cumate was added to a final concentration of 50 μM and cultures were grown at 30°C for 6 hours. Cultures were crosslinked with 1% (w/v) formaldehyde for 10 min, then crosslinking was quenched by addition of 125 mM glycine for 5 min. Cells were centrifuged at 7196 x g for 5 min at 4°C, supernatant was removed, and pellets were washed in 25 ml 1x cold PBS pH 7.5 three times. Pellets were resuspended in 1 ml [10 mM Tris pH 8 at 4°C, 1 mM EDTA, protease inhibitor tablet, 1 mg ml^-1^ lysozyme] and incubated at 37°C for 30 min. Sodium dodecyl sulfate (SDS) was added to a final concentration of 0.1% (w/v) and DNA was sheared to 300-500 bp fragments by sonication for 10 cycles (20 sec on/off). Debris was centrifuged at 15,000 x g for 10 min at 4°C, supernatant was transferred, and Triton X-100 was added to a final concentration of 1% (v/v). Samples were pre-cleared through incubation with 30 μl SureBeads Protein A magnetic beads for 30 min at room temp. Supernatant was transferred and 5% lysate was removed for use as input DNA.

For pulldown, 100 μl Pierce anti-FLAG magnetic agarose beads (25% slurry) were equilibrated overnight at 4°C in binding buffer [10 mM Tris pH 8 at 4°C, 1 mM EDTA, 0.1% (w/v) SDS, 1% (v/v) Triton X-100] supplemented with 1% (w/v) bovine serum albumin (BSA). Pre-equilibrated beads were washed four times in binding buffer, then incubated with the remaining lysate for 3 hours at room temperature. Beads were washed with low-salt buffer [50 mM HEPES pH 7.5, 1% (v/v) Triton X-100, 150 mM NaCl], high-salt buffer [50 mM HEPES pH 7.5, 1% (v/v) Triton X-100, 500 mM NaCl], and LiCl buffer [10 mM Tris pH 8 at 4°C, 1 mM EDTA, 1% (w/v) Triton X-100, 0.5% (v/v) IGEPAL CA-630, 150 mM LiCl]. To elute protein-DNA complexes, beads were incubated for 30 min at room temperature with 100 μl elution buffer [10 mM Tris pH 8 at 4°C, 1 mM EDTA, 1% (w/v) SDS, 100 ng μl^-1^ 3xFLAG peptide] twice. Elutions were supplemented with NaCl and RNase A to a final concentration of 300 mM and 100 μg ml^-1^, respectively, and incubated at 37°C for 30 min. Then, samples were supplemented with Proteinase K to a final concentration of 200 μg ml^-1^ and incubate overnight at 65°C to reverse crosslinks. Input and elutions were purified with the Zymo ChIP DNA Clean & Concentrator kit and libraries were prepared and sequenced at the Microbial Genome Sequencing Center (Pittsburgh, PA). Raw chromatin immunoprecipitation sequencing data are available in the NCBI GEO database under series accession GSE201499.

### ChIP-seq analysis

Paired-end reads were mapped to the *C. crescentus* NA1000 reference genome (GenBank accession number CP001340) with Bowtie2 on Galaxy. Peak calling was performed with the Genrich tool (https://github.com/jsh58/Genrich) on Galaxy; peaks are presented in Table S2. Briefly, PCR duplicates were removed from mapped reads, replicates were pooled, input reads were used as the control dataset, and peak were called using the default peak calling option [Maximum q-value: 0.05, Minimum area under the curve (AUC): 20, Minimum peak length: 0, Maximum distance between significant sites: 100]. An average AUC > 2500 was used as the cutoff for significant peaks. Distance between called peaks and the nearest transcription start sites (TSS) (modified from [32]) was analyzed using ChIPpeakAnno [66]. For genes/operons that did not have an annotated TSS, the +1 residue of the gene (or start of the operon) was designated as the TSS.

### RtrC motif discovery

For motif discovery, sequences of enriched ChIP-seq peaks were submitted to the XSTREME module of MEME suite [33]. For the XSTREME parameters, shuffled input files were used as the control sequences for the background model, checked for motifs between 6 and 30 bp in length that had zero or one occurrence per sequence.

### RNA preparation, sequencing, and analysis

Strains were incubated in quadruplicate at 30°C overnight in M2X broth supplemented with 50 μM cumate. Overnight replicate cultures were diluted into fresh M2X/50 μM cumate to 0.025 OD_660_ and incubated at 30°C for 8 hours. Cultures were diluted into 10 ml M2X/50 μM cumate to 0.001 – 0.003 OD_660_ and incubated at 30°C until reaching 0.3 – 0.4 OD_660_. Upon reaching the desired OD_660_, 6 ml culture was pelleted at 15,000 x g for 1 minute, supernatant was removed, and pellets were resuspended in 1 ml TRIzol and stored at -80°C. Samples were heated at 65°C for 10 min. Then, 200 μl chloroform was added, samples were vortexed, and incubated at room temperature for 5 min. Samples were centrifuged at 17,000 x g for 15 min at 4°C, then the upper aqueous phase was transferred to a fresh tube, an equal volume of 100% isopropanol was added, and samples were stored at -80°C overnight. Samples were centrifuged at 17,000 x g for 30 min at 4°C, then supernatant was removed. Samples were washed with cold 70% ethanol. Samples were centrifuged at 17,000 x g for 5 min at 4°C, supernatant was removed, and the pellet was allowed to dry. Pellets were resuspended in 100 μl RNase-free water and incubated at 60°C for 10 min. Samples were treated with TURBO DNase and cleaned up with RNeasy Mini Kit (Qiagen). Library preparation and sequencing was performed at the Microbial Genome Sequencing center with the Illumina Stranded RNA library preparation and RiboZero Plus rRNA depletion (Pittsburgh, PA). Reads were mapped to the *C. crescentus* NA1000 reference genome (GenBank accession number CP001340) using CLC Genomics Workbench 20 (Qiagen). Differential gene expression was determined with the CLC Genomics Workbench RNA-seq Analysis Tool (|fold-change| > 1.5 and FDR p-value < 0.001). Raw RNA sequencing data are available in the NCBI GEO database under series accession GSE201499.

### RNA-seq and ChIP-seq overlap analysis

The Bioconductor package was used to identify overlap between RtrC-regulated genes defined by RNA-seq and RtrC binding sites defined by ChIP-seq [66]. Promoters for genes were designated as the sequence -400 to +100 around the TSS [32]. Overlap between promoters and RtrC motifs identified from XSTREME was analyzed using ChIPpeakAnno within Bioconductor. Genes were defined as direct targets of RtrC if their transcript levels were differentially regulated in the RNA-seq analysis and had an RtrC motif within a promoter for their operon. To analyze RtrC motif distribution in directly regulated promoters, promoters were grouped based on the effect of RtrC on gene expression (i.e. upregulated vs. downregulated). The number of predicted RtrC motifs at each position with these promoters was calculated and plotted. If an operon contained more than one promoter, then each promoter for that operon that contained an RtrC motif was analyzed.

### Analysis of transcription using fluorescent fusions

Strains were incubated in triplicate at 30°C overnight in PYE or M2X broth supplemented with 1 μg ml^-1^ chloramphenicol and 50 μM cumate. Overnight cultures were diluted to 0.05 OD_660_ in the appropriate broth and incubated at 30°C for 24 hours. For Figure S4 and S5, log phase (0.05 – 0.3 OD_660_) cultures were diluted to 0.025 OD_660_ and incubated at 30°C for 24-48 hours. Then, 200 μl culture was transferred to a black Costar 96 well plate with clear bottom (Corning). Absorbance at 660 nm and fluorescence (excitation = 497 + 10 nm; emission = 523 + 10 nm) were measured in a Tecan Spark 20M plate reader. Fluorescence was normalized to absorbance.

For Figure 6, strains were incubated in triplicate at 30°C overnight in PYE broth supplemented with 1 μg ml^-1^ chloramphenicol. Overnight cultures were diluted to 0.025 OD_660_ in PYE broth supplemented with 1 μg ml^-1^ chloramphenicol, 50 μM cumate, and 0.15% (w/v) xylose. Cultures were incubated at 30°C for 24 hours, then 100 μl overnight was diluted with 100 μl PYE and transferred to a black Costar 96 well plate with clear bottom (Corning). Fluorescence and absorbance were measured as indicated above in a Tecan Spark 20M plate reader. Fluorescence was normalized to absorbance. Statistical analysis was carried out in GraphPad 9.3.1.

### Protein purification

For heterologous expression of RtrC, plasmids were transformed into the BL21 Rosetta (DE3)/pLysS background. Strains were inoculated into 20 ml LB broth supplemented with 100 μg ml^-1^ carbenicillin and incubated overnight at 37°C. Overnight cultures were diluted into 1000 ml LB supplemented with carbenicillin and grown for 3 – 4 hours at 37°C. Protein expression was induced by 0.5 mM isopropyl β-D-1-thiogalactopyranoside (IPTG) and incubation at 37°C for 3 – 4.5 hours. Cells were pelleted at 11,000 x g for 7 min at 4°C, pellets were resuspended in 25 ml lysis buffer [25 mM Tris pH 8 at 4°C, 500 mM NaCl, 10 mM imidazole], and stored at -80°C. Samples were thawed, supplemented with PMSF and benzonase to a final concentration of 1 mM and 50 Units ml^-1^, respectively. Samples were sonicated with a Branson Digital Sonifier at 20% output in 20” intervals until sufficiently lysed and clarified by centrifugation at 39,000 x g for 15 min at 4°C. Clarified lysates were batch incubated with 4 ml Ni-NTA Superflow Resin (50% slurry) that had been pre-equilibrated in lysis buffer for 60 min at 4°C. Column was then washed with 25 ml lysis buffer, high salt buffer [25 mM Tris pH 8 at 4°C, 1 M NaCl, 30 mM imidazole], and low salt buffer [25 mM Tris pH 8 at 4°C, 500 mM NaCl, 30 mM imidazole]. For elution, column was batch incubated with 25 ml elution buffer [25 mM Tris pH 8 at 4°C, 500 mM NaCl, 300 mM imidazole] for 60 min at 4°C.

Elution was supplemented with ULP1 protease to cleave the His_6_-SUMO tag and dialyzed in 1 L dialysis buffer [25 mM Tris pH 8 at 4°C, 500 mM NaCl] overnight at 4°C. Dialyzed sample was batch incubated with 4 ml Ni-NTA Superflow Resin (50% slurry) that had been pre-equilibrated in dialysis buffer for 60 min at 4°C. Flowthrough that contained untagged RtrC was collected and concentrated on an Amicon Ultra-15 concentrator (3 kDa cutoff) at 4,000 x g at 4°C. Samples were stored at 4°C until needed.

### Electrophoretic mobility shift assay (EMSA)

To prepare labeled DNA fragments, an Alexa488-labeled universal forward primer and an *hfiA* specific reverse primer listed in Table S5 were annealed in a thermocycler in as follows: 94°C for 5 min, then ramp down to 18°C at 0.1°C s^-1^. Overhangs were filled in with DNA polymerase I, Large Klenow fragment at 25°C for 60 min. DNA fragments were then treated with Mung Bean Nuclease for 120 min at 30°C to remove any remaining overhangs. DNA fragments were purified with the GeneJet PCR purification kit, eluted in 10 mM TE/NaCl [Tris pH 8 at 4°C, 1 mM EDTA, 50 mM NaCl], and diluted to 0.5 μM in TE/NaCl. Unlabeled DNA fragments were prepared by annealing primers listed in Table S5 with protocol listed above. For non-specific chase, the sequence of the *hfiA* specific probe was shuffled.

RtrC was incubated with 6.25 nM labeled DNA in binding buffer at 20°C for 30 min in the dark and subsequently cooled to 4°C on ice. For EMSA to analyze binding curves, DNA binding buffer consisted of 32.5 mM Tris pH 8 at 4°C, 200 mM NaCl, 1 mM EDTA, 30% (v/v) glycerol, 1 mM DTT, 10 μg ml^-1^ BSA, and 50 ng μl^-1^ poly(dI-dC). For EMSAs with unlabeled chases, DNA binding buffer consisted of 30 mM Tris pH 8 at 4°C, 150 mM NaCl, 1 mM EDTA, 30% (v/v) glycerol, 1 mM DTT, 10 μg ml^-1^ BSA, and 50 ng μl^-1^ poly(dI-dC). For non-specific and specific chases, reactions were supplemented with 2.5 μM unlabeled DNA. Then, 15 μl reaction was loaded on to a degassed polyacrylamide gel [10% (v/v/) acrylamide (37.5:1 acrylamide:bis-acrylamide), 0.5x Tris-Borate-EDTA buffer (TBE: 45 mM Tris, 45 mM borate, 1 mM EDTA)] and run at 100 V for 40 min at 4°C in 0.5x TBE buffer. Gels were imaged on a ChemiDoc MP imaging system [Light: blue epi illumination, Filter: 530/28, Exposure: 30 sec] and bands were quantified with FIJI. For calculating k_d_, percent bound probe at each protein concentration was calculated as (1 – [intensity free probe at x nM protein]/[intensity of free probe at 0 nM protein]). Binding curve was derived from One site – specific binding analysis using GraphPad 9.3.1.

### Tn-himar-seq to assess gene contributions to fitness in a pellicle biofilm

We grew a barcoded Tn-himar mutant library previously described and characterized [38] in static culture and harvested cells from the air-liquid interface using an approach described in [37]. For the experiment in complex (PYE) medium, a 1 ml aliquot of the library was diluted into 100 ml of PYE in a 250 ml flask and outgrown shaking at 30°C overnight. Five aliquots of 200 μl of the starter culture were saved as a reference sample. Five beakers, each containing 400 ml PYE, were inoculated to a starting density of OD_660_ = 0.005-0.006. These beakers were incubated at room temperature without shaking and samples from the air-liquid interface were collected using the large end of sterile 1 ml pipet tips [37]. The interfacial liquid and cells collected in the pipet tip were transferred to a 1.7 ml centrifuge tube containing 1.5 ml of sterile water. Contact with the sterile water allowed efficient transfer of the interfacial sample to the tube. At early time points (less than 2 days), two sample plugs were collected from the interface of each replicate beaker. At later time points (2+ days), one sample plug contained sufficient numbers of cells for analysis. After transfer, cells were collected by centrifugation (3 min at 21,000g), the supernatant was removed, and the cell pellet was stored at -20°C. The experiment in defined M2X medium, was conducted similarly, except that the starter culture was grown in M2X medium and samples were collected on a slower and longer time course because pellicles develop more slowly in defined medium [37].

To assess barcode abundances, we followed the approach developed and described by Wetmore and colleagues [61]. Briefly, each cell pellet was resuspended in 10-20 μl water. Barcodes were amplified using Q5 polymerase (New England Biolabs) in 20 μl reaction volumes containing 1X Q5 reaction buffer, 1X GC enhancer, 0.8 U Q5 polymerase, 0.2 mM dNTP, 0.5 μM of each primer and 1 μl of resuspended cells. Each reaction contained a universal forward primer, Barseq_P1, and a unique indexed reverse primer (Barseq_P2_ITxxx, where the xxx identifies the index number) described in [61]. Reactions were cycled as follows: 98 °C for 4 min, 25 cycles of 98 °C for 30 s, 55 °C for 30 s, and 72 °C for 30, 72 °C for 5 min, 4°C hold. Amplified barcodes were pooled and 50-bp single-end reads were collected on an Illumina HiSeq4000 with Illumina TruSeq primers at the University of Chicago Genomics Facility. Pellicle barcode amplicon sequence data have been deposited in the NCBI Sequence Read Archive under BioProject accession PRJNA877623. Sequence data used to map the Tn insertion sites to the *Caulobacter crescentus* genome are available under BioProject accession PRJNA429486, SRA accession SRX3549727.

Barcode sequences were analyzed using the fitness calculation protocol of Wetmore and colleagues [61]. Briefly, the barcodes in each sample were counted and assembled using MultiCodes.pl and combineBarSeq.pl. From this table of barcodes, FEBA.R was used to determine fitness by comparing the log_2_ ratios of barcode counts in each sample over the counts in the starter culture reference samples. Fitness scores corresponding to the genes of interest in this study were manually extracted.

### Soft agar swarm assay

Strains were incubated in quadruplicate at 30°C overnight in PYE broth supplemented with 50 μM cumate. Overnight cultures were diluted to 0.05 OD_660_ in PYE/50 μM cumate, then incubated at 30°C for 24 hours. Cultures were diluted to 0.5 OD_660_ in PYE broth, 0.75 μl diluted culture was pipetted into PYE plate supplemented with 50 μM cumate, incubated at 30°C for 3 days. Plates were imaged on a ChemiDoc MP imaging system and swarm size was measured with FIJI. Statistical analysis was carried out in GraphPad 9.3.1.

### Neighborhood analysis

RtrC protein sequence was compared to the NCBI Refseq database with PSI-BLAST, using the default settings and excluding uncultured/environmental samples. Accession ID for proteins with > 95% query coverage and > 65% percent identity were extracted and submitted to the WebFLaGs server for neighborhood analysis (http://www.webflags.se/) [67].

## Supplementary Figure Legends

**Figure S1. *hfiA* promoter architecture**. Schematic of the *hfiA* promoter. Binding sites for CtrA, StaR and RtrC are marked with pink, grey, and blue boxes, respectively. Experimentally mapped transcription start sites are marked with black arrows. The start of the *hfiA* coding region is marked with an orange box. Sites previously identified in a screen for mutations that result in increased expression from the *hfiA* promoter [16] are marked by dark grey boxes with the corresponding mutations (white lettering).

**Figure S2. RtrC binds DNA *in vitro*. A)** Electrophoretic mobility shift assay (EMSA) using purified RtrC. Increasing concentrations of purified RtrC (0, 12.5, 25, 50, 75, 100, 150, and 300 nM) were incubated with 6.25 nM labeled *hfiA* probe. Blot is representative of three biological replicates. **B)** RtrC DNA binding curve derived from triplicate EMSA data. K_d_ was calculated based on assumption of one site specific binding.

**Figure S3. Genes that contain an RtrC motif in their promoters but that are not differentially regulated by *rtrC* overexpression**. There are several genes that are not regulated by *rtrC* overexpression in the RNA-seq dataset (Table S3) despite the presence of an RtrC binding site in their promoter. To confirm this result, transcription from these genes was measured using P_*gene*_-*mNeonGreen* transcriptional fusion reporters in an empty vector (EV) or *rtrC* overexpression strain (*rtrC*^++^). Cells grown in complex medium (PYE) and fluorescence was normalized to cell density (OD_660_). Data show the mean; error bars represent standard deviation of three biological replicates. Statistical significance was determined by multiple unpaired t tests, correcting for multiple comparisons using the Holm-Šídák method (ns – not significant).

**Figure S4. *rtrB* expression in logarithmic versus stationary phase**. *rtrB* transcription can be activated by both SpdR and RtrC (see Figure 6). *rtrB* transcription was measured using a P_*rtrB*_-*mNeonGreen* transcriptional fusion reporter in a wild type (WT) or *spdR*_D64E_ background with in-frame deletions (Δ) in *spdR* and/or *rtrC*. Cells were grown in complex medium (PYE) to early logarithmic phase (marked as 0 h) and cultivated for an additional 48 h into stationary phase (marked as 48 h); fluorescence was measured at the 0 h and 48 h points (see methods). Fluorescence measurements were normalized to cell density (OD_660_). Data are the mean; errors bars represent standard deviation of three biological replicates. Statistical significance was determined by Two-way ANOVA followed by Tukey’s multiple comparisons within each time point (p-value < 0.01,**; p-value < 0.0001,****; ns – not significant).

**Figure S5. *rtrA, rtrB, and rtrC* expression is regulated in a media-, growth phase- and *spdR*-dependent manner**. *rtrA, rtrB*, and *rtrC* transcription was measured with P_*rtrA*_-*mNeonGreen* (*mNG*), P_*rtrB*_-*mNG*, and P_*rtrC*_-*mNG* reporters, respectively, in wild type (WT) or a strain bearing an in-frame deletion (Δ) in *spdR*. Cells were grown in complex (PYE) or defined medium (M2-xylose) to early logarithmic phase and measured (0 h) or to stationary phase (24 h). Fluorescence was normalized to cell density (OD_660_). Data are the mean; errors bars represent standard deviation of three biological replicates. Statistical significance was determined by two-way ANOVA followed by Šídák multiple comparison test (p-value < 0.001,***; p-value < 0.0001,****; ns – not significant).

**Figure S6. Regulation of holdfast synthesis in complex media as a function of growth phase. A-B)** Percentage of cells with stained holdfast in wild type (WT) and in strains bearing in-frame deletions (Δ) of *spdR, rtrA, rtrB*, or *rtrC*, or an *rtrABC* triple deletion. Holdfast counts were performed on cultures grown in complex medium (PYE) in **A)** early log phase or **B)** stationary phase (after 24 hours of growth). Data show the mean holdfast percentage; error bars are standard deviation of three biological replicates. Statistical significance was determined by one-way ANOVA followed by Dunnett’s multiple comparison (p-value < 0.001,***).

**Figure S7. Transposon insertions holdfast synthesis (*hfsA*), in adhesion TCS genes, and in *rtrC* affect temporal adhesion profiles in cheesecloth and pellicle assays. A)** Fitness timecourse of *C. crescentus* mutants harboring transposon insertions in adhesion regulators (*lovK, spdS, skaH, spdR*, and *rtrC*) and a representative holdfast synthesis gene (*hfsA*) over 5 days of serial passaging in the presence of cheesecloth. Strains were sampled from the supernatant, outside of the cheesecloth, which is enriched with non-adherent cells [22] (n=3) **B)** Fitness timecourse of *C. crescentus* mutants harboring transposon insertions in adhesion regulators and a representative holdfast synthesis gene sampled from a pellicle biofilm over 4 days of static cultivation in complex (PYE) medium (n=5) **C)** Fitness timecourse of *C. crescentus* mutants harboring transposon insertions in adhesion regulators and a representative holdfast synthesis gene sampled from a pellicle biofilm over 7 days of static cultivation in M2-xylose defined medium (n=4). Data show the mean; errors bars represent standard deviation of at least three biological replicates.

**Figure S8. *rtrC* genomic neighborhood is conserved in *Caulobacter***. Phylogenetic tree based on RtrC sequence (left) and genomic neighborhood (right) surrounding *rtrC* in various bacterial species. Protein sequence accessions were retrieved from the NCBI RefSeq database by a PSI-BLAST search and analyzed with the webFLaGs server (http://www.webflags.se/) [67]. Numbers on phylogenetic tree indicate bootstrap values. *rtrC* homologs are colored black, orthologous genes are colored and numbered identically, non-conserved genes are uncolored and outlined in grey, pseudogenes are uncolored and outlined in blue, and non-coding RNA genes are colored green.

