## Supplemental Figures for "A cryptic transcription factor regulates *Caulobacter* adhesin development"

**Supplementary Figures**

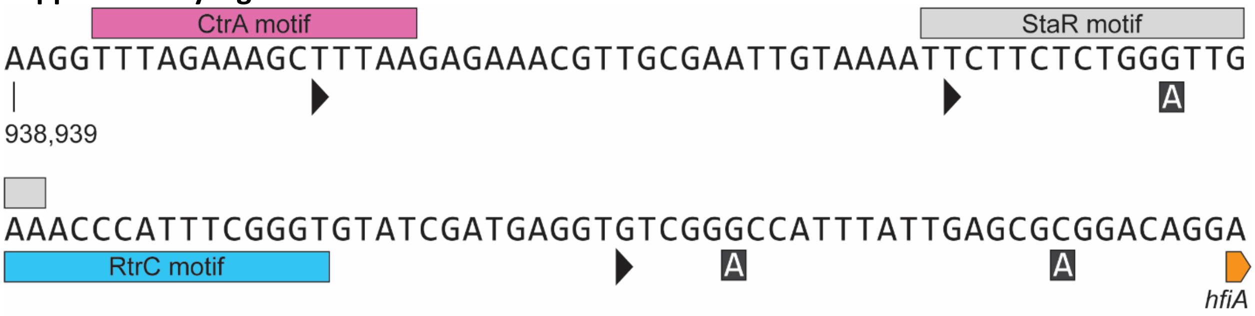


**Figure S1. *hfiA* promoter architecture.** Schematic of the *hfiA* promoter. Binding sites for CtrA, StaR and RtrC are marked with pink, grey, and blue boxes, respectively. Experimentally mapped transcription start sites are marked with black arrows. The start of the *hfiA* coding region is marked with an orange box. Sites previously identified in a screen for mutations that result in increased expression from the *hfiA* promoter [16] are marked by dark grey boxes with the corresponding mutations (white lettering).

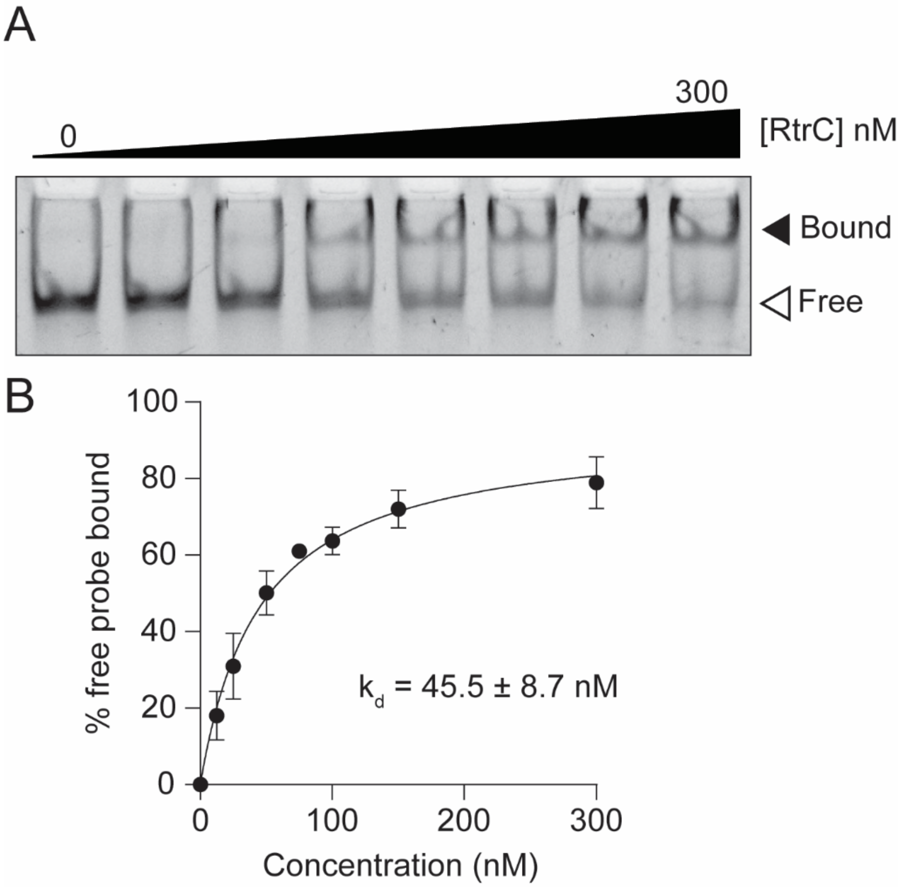


**Figure S2. RtrC binds DNA *in vitro*. A)** Electrophoretic mobility shift assay (EMSA) using purified RtrC. Increasing concentrations of purified RtrC (0, 12.5, 25, 50, 75, 100, 150, and 300 nM) were incubated with 6.25 nM labeled *hfiA* probe. Blot is representative of three biological replicates. **B)** RtrC DNA binding curve derived from triplicate EMSA data. $K_d$ was calculated based on assumption of one site specific binding.

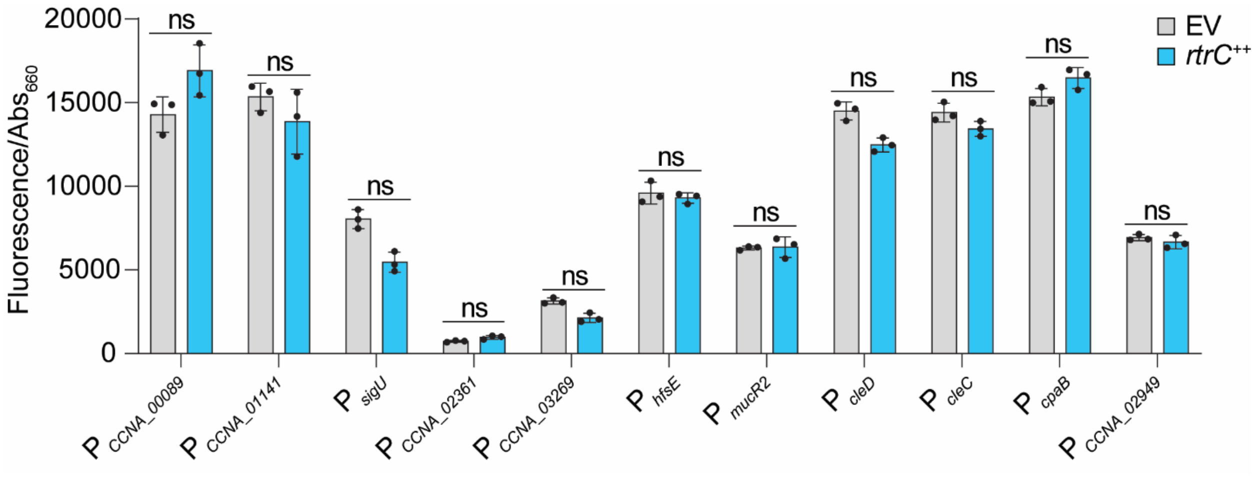


**Figure S3. Genes that contain an RtrC motif in their promoters but that are not differentially regulated by *rtrC* overexpression.** There are several genes that are not regulated by *rtrC* overexpression in the RNA-seq dataset (Table S3) despite the presence of an RtrC binding site in their promoter**.** To confirm this result, transcription from these genes was measured using $P_{gene}$-*mNeonGreen* transcriptional fusion reporters in an empty vector (EV) or *rtrC* overexpression strain (*rtrC*$^{++}$). Cells grown in complex medium (PYE) and fluorescence was normalized to cell density ($OD_{660}$). Data show the mean; error bars represent standard deviation of three biological replicates. Statistical significance was determined by multiple unpaired t tests, correcting for multiple comparisons using the Holm-Šídák method (ns – not significant).

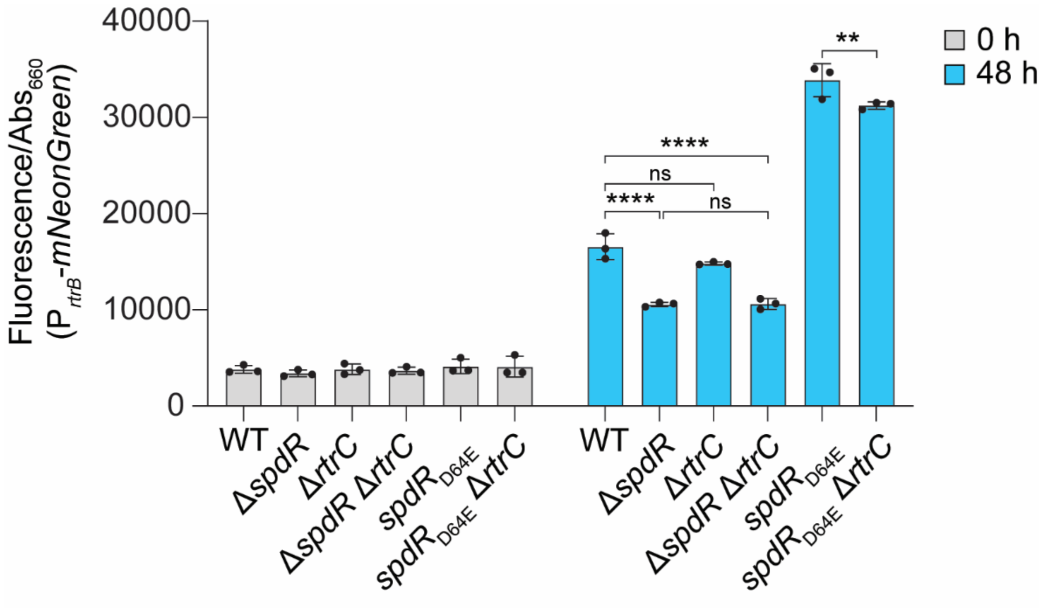


**Figure S4. *rtrB* expression in logarithmic versus stationary phase.** *rtrB* transcription can be activated by both SpdR and RtrC (see Figure 6). *rtrB* transcription was measured using a $P_{rtrB}$-*mNeonGreen* transcriptional fusion reporter in a wild type (WT) or *spdR*$_{D64E}$ background with in-frame deletions (Δ) in *spdR* and/or *rtrC*. Cells were grown in complex medium (PYE) to early logarithmic phase (marked as 0 h) and cultivated for an additional 48 h into stationary phase (marked as 48 h); fluorescence was measured at the 0 h and 48 h points (see methods). Fluorescence measurements were normalized to cell density ($OD_{660}$). Data are the mean; errors bars represent standard deviation of three biological replicates. Statistical significance was determined by Two-way ANOVA followed by Tukey's multiple comparisons within each time point (p-value $\leq$ 0.01,**; p-value $\leq$ 0.0001,****; ns – not significant).

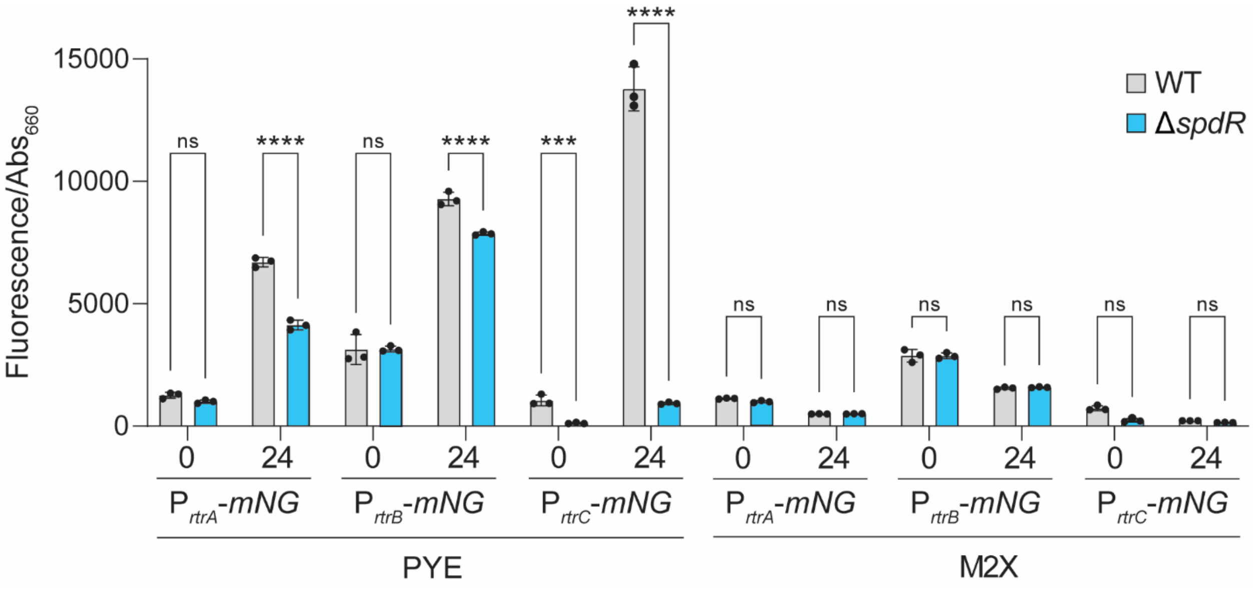


**Figure S5.** ***rtrA, rtrB, and rtrC*** **expression is regulated in a media-, growth phase- and** ***spdR*****-dependent manner.** *rtrA*, *rtrB*, and *rtrC* transcription was measured with $P_{rtrA}$-*mNeonGreen* (*mNG*), $P_{rtrB}$-*mNG*, and $P_{rtrC}$-*mNG* reporters, respectively, in wild type (WT) or a strain bearing an in-frame deletion (Δ) in *spdR*. Cells were grown in complex (PYE) or defined medium (M2-xylose) to early logarithmic phase and measured (0 h) or to stationary phase (24 h). Fluorescence was normalized to cell density ($OD_{660}$). Data are the mean; errors bars represent standard deviation of three biological replicates. Statistical significance was determined by two-way ANOVA followed by Šídák multiple comparison test (p-value $\leq$ 0.001,***; p-value $\leq$ 0.0001,****; ns – not significant).

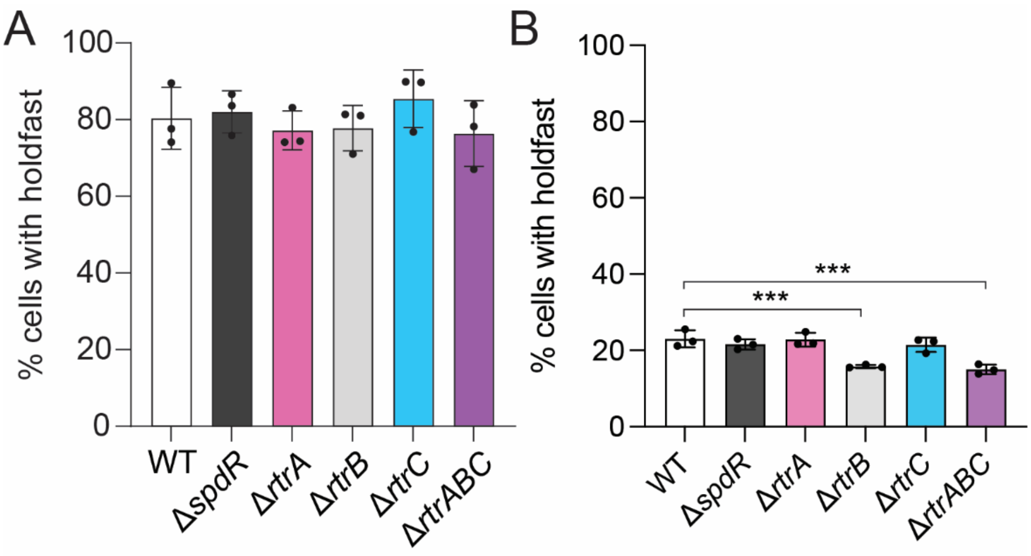


**Figure S6. Regulation of holdfast synthesis in complex media as a function of growth phase.**

**A-B)** Percentage of cells with stained holdfast in wild type (WT) and in strains bearing in-frame deletions (Δ) of *spdR*, *rtrA*, *rtrB*, or *rtrC*, or an *rtrABC* triple deletion. Holdfast counts were performed on cultures grown in complex medium (PYE) in **A)** early log phase or **B)** stationary phase (after 24 hours of growth). Data show the mean holdfast percentage; error bars are standard deviation of three biological replicates. Statistical significance was determined by one-way ANOVA followed by Dunnett's multiple comparison (p-value $\leq$ 0.001,***).

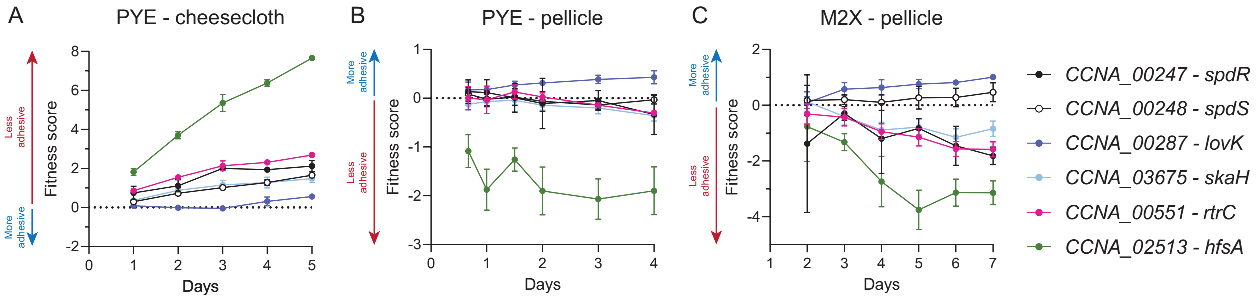


**Figure S7. Transposon insertions holdfast synthesis (*hfsA*), in adhesion TCS genes, and in *rtrC* affect temporal adhesion profiles in cheesecloth and pellicle assays. A)** Fitness timecourse of *C. crescentus* mutants harboring transposon insertions in adhesion regulators (*lovK, spdS, skaH, spdR*, and *rtrC*) and a representative holdfast synthesis gene (*hfsA*) over 5 days of serial passaging in the presence of cheesecloth. Strains were sampled from the supernatant, outside of the cheesecloth, which is enriched with non-adherent cells [22] (n=3) **B)** Fitness timecourse of *C. crescentus* mutants harboring transposon insertions in adhesion regulators and a representative holdfast synthesis gene sampled from a pellicle biofilm over 4 days of static cultivation in complex (PYE) medium (n=5) **C)** Fitness timecourse of *C. crescentus* mutants harboring transposon insertions in adhesion regulators and a representative holdfast synthesis gene sampled from a pellicle biofilm over 7 days of static cultivation in M2-xylose defined medium (n=4). Data show the mean; errors bars represent standard deviation of at least three biological replicates.

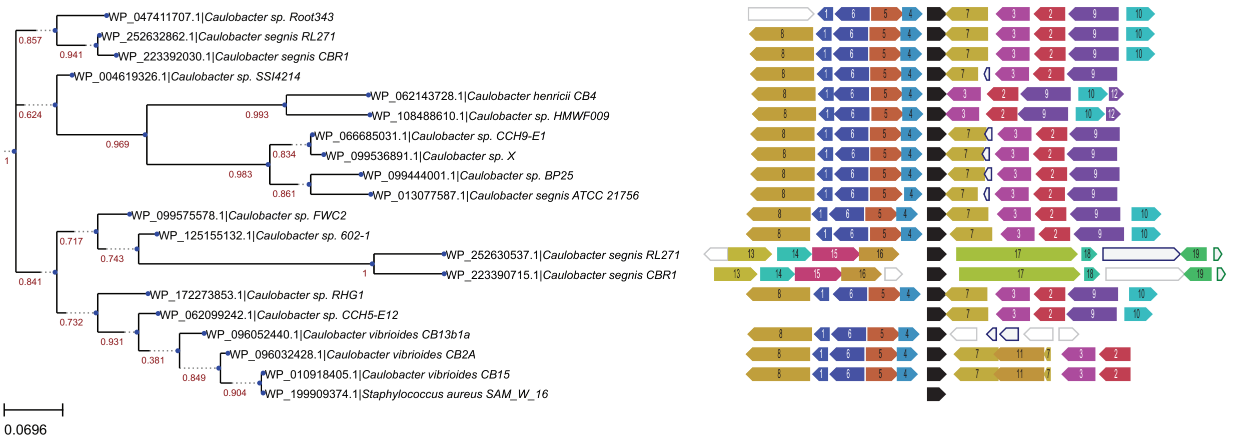


**Figure S8. *rtrC* genomic neighborhood is conserved in *Caulobacter*.** Phylogenetic tree based on RtrC sequence (left) and genomic neighborhood (right) surrounding *rtrC* in various bacterial species. Protein sequence accessions were retrieved from the NCBI RefSeq database by a PSI-BLAST search and analyzed with the webFLaGs server (http://www.webflags.se/) [67]. Numbers on phylogenetic tree indicate bootstrap values. *rtrC* homologs are colored black, orthologous genes are colored and numbered identically, non-conserved genes are uncolored and outlined in grey, pseudogenes are uncolored and outlined in blue, and non-coding RNA genes are colored green.
